# Fast-spiking interneuron detonation drives high-fidelity inhibition in the olfactory bulb

**DOI:** 10.1101/2024.05.07.592874

**Authors:** Shawn D. Burton, Christina M. Malyshko, Nathaniel N. Urban

**Affiliations:** Department of Biological Sciences, Lehigh University, Bethlehem, PA, USA

## Abstract

Inhibitory circuits in the mammalian olfactory bulb (OB) dynamically reformat olfactory information as it propagates from peripheral receptors to downstream cortex. To gain mechanistic insight into how specific OB interneuron types support this sensory processing, we examine unitary synaptic interactions between excitatory mitral and tufted cells (MTCs), the OB projection cells, and a conserved population of anaxonic external plexiform layer interneurons (EPL-INs) using pair and quartet whole-cell recordings in acute mouse brain slices. Physiological, morphological, neurochemical, and synaptic analyses divide EPL-INs into distinct subtypes and reveal that parvalbumin-expressing fast-spiking EPL-INs (FSIs) perisomatically innervate MTCs with release-competent dendrites and synaptically detonate to mediate fast, short-latency recurrent and lateral inhibition. Sparse MTC synchronization supralinearly increases this high-fidelity inhibition, while sensory afferent activation combined with single-cell silencing reveals that individual FSIs account for a substantial fraction of total network-driven MTC lateral inhibition. OB output is thus powerfully shaped by detonation-driven high-fidelity perisomatic inhibition.

## Introduction

Circuit operations underlying mammalian brain function critically depend on the diverse structural and functional features of distinct inhibitory interneuron types, such as the dynamic modulation of projection cell output gain, spike time patterning, and synchronization by fast-spiking perisomatic-innervating basket cells in neocortex and hippocampus (Tiesinga et al., 2008; Silver, 2010; Hu et al., 2014). Resolving the functions of interneurons in these circuits through cell type-specific recordings of unitary synaptic interactions has dramatically advanced our understanding of brain function across learning and disease (Marín, 2012). In the olfactory bulb (OB), the first processing station of the main olfactory system, inhibitory interneurons likewise exhibit pronounced diversity (Nagayama et al., 2014; Burton, 2017) and are similarly central to OB circuit operations, with perturbations of synaptic inhibition significantly disrupting projection mitral/tufted cell (MTC) spike time patterning and olfactory-guided behavior (Fukunaga et al., 2012; Lepousez and Lledo, 2013; Gödde et al., 2016). In contrast to the extensive progress made in linking diverse interneuron types to specific functions elsewhere in the brain, however, understanding how specific inhibitory interneurons support OB circuit operations remains limited.

Conceptual models of how synaptic inhibition shapes OB activity focus predominantly on granule cells (GCs), the most abundant type of OB interneuron (Shepherd et al., 2004; Burton, 2017). Despite theoretical and histological evidence of widespread dendrodendritic MTC–GC connections (Rall et al., 1966; Rall and Shepherd, 1968), functional unitary MTC–GC connectivity appears exceedingly sparse, with <5% of pair recordings exhibiting MTC-to-GC excitation and *zero* exhibiting GC-to-MTC inhibition (Isaacson, 2001; Kato et al., 2013; Pressler and Strowbridge, 2017). Investigation of MTC inhibition has consequently relied on non-cell-type-specific measures of total recurrent inhibition evoked by prolonged MTC activation, assumed to originate from GCs, and often performed in low Mg^2+^ to augment GC excitation (Schoppa and Urban, 2003; Burton, 2017). These measures have led to the consensus that MTC inhibition is slow and low-fidelity (Galán et al., 2006; Kapoor and Urban, 2006), features difficult to reconcile with functions such as the precise regulation of MTC spike timing.

Fundamental understanding of the circuit operations underlying olfaction will thus ultimately require greater cell type-specific knowledge of unitary synaptic interactions in the OB. Of paramount interest, the external plexiform layer (EPL) contains a conserved and neurochemically diverse population of anaxonic interneurons (EPL-INs) that, while less abundant than GCs, can mediate unitary MTC inhibition (Huang et al., 2013; Kato et al., 2013). Population-level manipulations further suggest that such inhibition powerfully influences MTC activity (Huang et al., 2013; Kato et al., 2013; Liu et al., 2019; Wang et al., 2022). However, interpretation of these genetically-targeted approaches remains constrained by disruption of endogenous expression in at least some mouse lines (Viollet et al., 2017; Joye et al., 2020; Wang et al., 2022), nonselective perturbation of other neuron types (Kato et al., 2013; Wang et al., 2021; 2022), incomplete mapping of neurochemical identity across EPL-IN subtypes, and limited mechanistic insight into unitary interactions, leaving the overall function of EPL-INs still unclear.

To advance our fundamental understanding of circuit operations in the OB, we have therefore performed a systematic investigation of unitary synaptic interactions between MTCs and EPL-INs using simultaneous acute-slice whole-cell recordings in physiological Mg^2+^. Our results reveal that fast-spiking EPL-INs perisomatically innervate MTCs with noncanonical architecture and mediate a substantial fraction of total MTC inhibition through fast, synchronous synaptic release. This release, supporting high-fidelity recurrent and lateral inhibition, is driven by both unusually prevalent synaptic detonation and high sensitivity to sparse MTC synchronization. Collectively, these findings challenge multiple conceptual paradigms of OB circuit operation and provide new insight into key modes of inhibitory signaling supporting olfaction.

## Results

### EPL-INs comprise two subtypes with distinct circuit functions

To systematically investigate unitary MTC–EPL-IN interactions independent of neurochemical identity and molecular lineage, we targeted MTCs and nearby small somata of putative EPL-INs for whole-cell pair recordings in acute OB slices prepared from mice devoid of genetically-targeted interneuronal labeling. Despite wide neurochemical heterogeneity noted across EPL-INs subsets (e.g., Lepousez et al., 2010; Huang et al., 2013), analysis of intrinsic biophysical properties across 145 MTC–EPL-IN pairs surprisingly revealed only two major EPL-IN subtypes: 1) fast-spiking interneurons (FSIs) with non-adapting spike trains, high instantaneous firing rates, and frequent spike clustering; and 2) regular-spiking interneurons (RSIs) with regular, adapting, and comparatively slower firing (Figure 1A-H). Unbiased hierarchical clustering (Figure 1I) and principal component analysis (Figure S1) of 23 intrinsic biophysical properties directly supported this classification, with FSIs and RSIs exhibiting stark differences across the majority of properties (Figure S2; Table S1).

**Figure 1.**
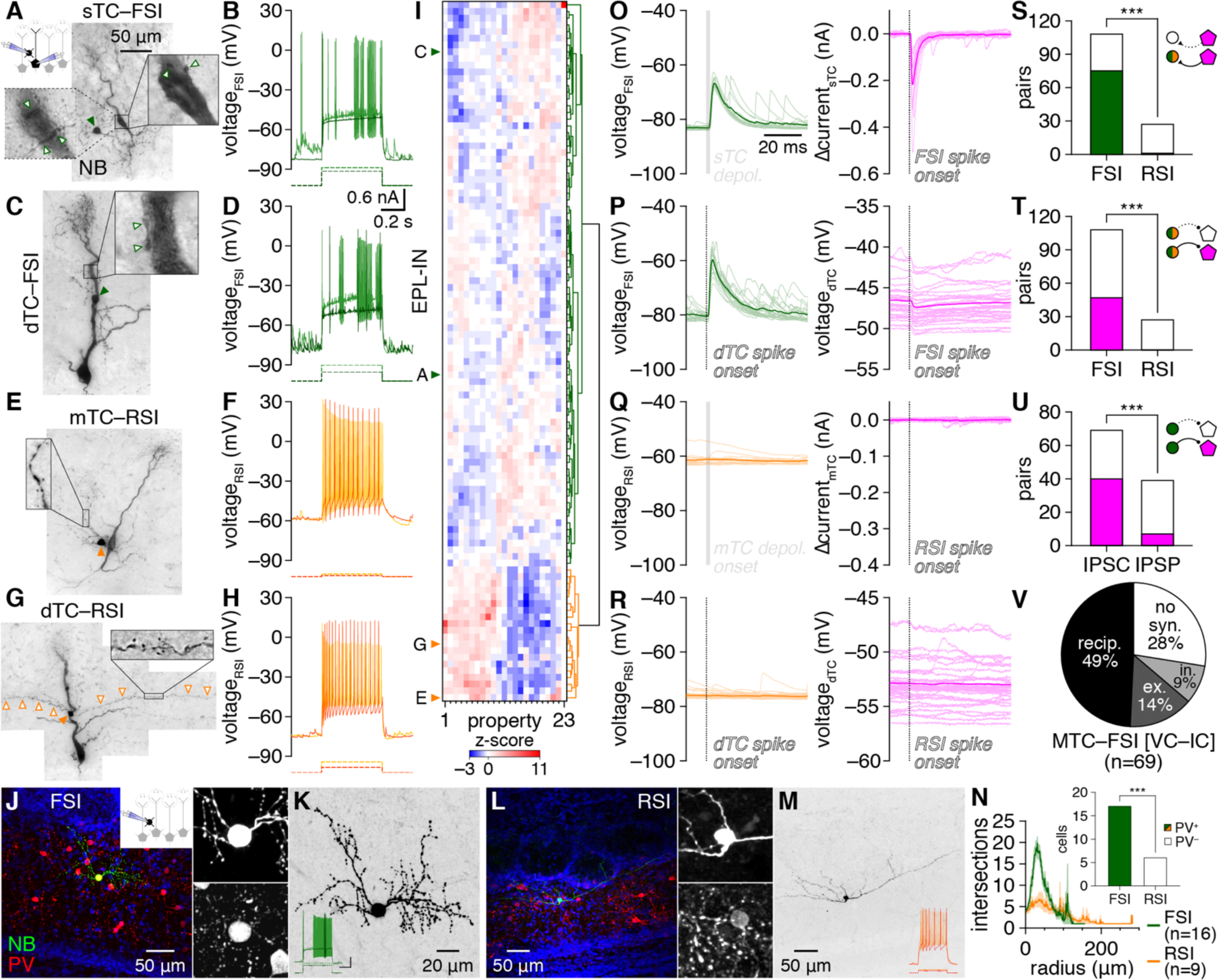
EPL-INs subdivide into fast- and regular-spiking subtypes. **A-D:** Example MTC–FSI pairs filled with NB (**A,C**) and interneuron fast-spiking physiology (**B,D**) in response to step current injections (dashed lines). Magnification here and throughout other figures shows single optical planes of example putative MTC–FSI synaptic contact (open arrowheads), except where noted. Schematic of recording configuration also shown here and throughout other figures when configuration changes. **E-H:** Example MTC– RSI pairs and regular-spiking physiology. Magnification: long spiny RSI dendrites (open arrowheads). Filled arrowheads: interneuron somata. **I:** Hierarchical clustering of 104 EPL-INs by intrinsic biophysical properties into FSIs (green) and RSIs (orange). **J:** Post hoc PV staining of an example FSI. Insets: soma magnification of NB (upper) and PV (lower). **K:** NB signal from **J**, inverted to show morphology. Inset: spiking physiology (scale: 20 mV/0.5 nA, 0.2 s). **L,M:** Same as **J,K** for an RSI. **N:** Sholl analysis. Inset: FSIs were PV^+^, while RSIs were PV^−^ (***p=1.6×10^−6^, χ^2^_[1]_=23, χ^2^ test). **O-R:** Unitary synaptic interactions from the pairs in **A-H**. **S,T:** Distribution of excitatory (**S**) (***p=7.3×10^−10^, χ^2^_[1]_=37.9, χ^2^ test) and inhibitory (**T**) (***p=2.2×10^−5^, χ^2^_[1]_=18.0, χ^2^ test) unitary connections among MTC–FSI and MTC–RSI pairs. **U:** Voltage-clamp (VC) detected unitary inhibitory connections more sensitively than current-clamp (IC) (***p=5.6×10^−5^, χ^2^_[1]_=16.2, χ^2^ test). **V:** Distribution of inhibition, excitatory, reciprocal, or no unitary connectivity within the subset of MTC–FSI pairs recorded in VC and IC, respectively.

FSIs and RSIs also exhibited morphological differences: FSIs extended complex and beaded dendritic arbors while RSIs extended less branched and long, spiny dendrites that occasionally entered the glomerular layer (Figure 1A,C,E,G,J-N; Figure S3; Figure S4; Figure S5; Table S2). Neither FSIs nor RSIs extended visible axons. Morphological properties of FSIs thus broadly matched those previously noted for several neurochemical EPL-IN subsets (Lepousez et al., 2010; Crespo et al., 2013; Huang et al., 2013; Kato et al., 2013), while RSIs did not obviously correspond to any previously-characterized EPL-INs. Consistent with these parallels, post hoc immunostaining confirmed parvalbumin (PV) expression in 100% of FSIs and 0% of RSIs (Figure 1N; Figure S3). Likewise, neither FSIs nor RSIs expressed tyrosine hydroxylase (TH), a marker of sparse, tonically-active short-axon cells in the EPL (Figure S4) (Pignatelli et al., 2009; Liberia et al., 2012; Kosaka et al., 2020). How other neurochemicals map to FSIs vs. RSIs remains unclear, however, with only 27% of FSIs exhibiting weakly-detectable vasoactive intestinal peptide (VIP) expression compared to 0% of RSIs (Figure S5).

Physiological EPL-IN differences also mapped directly onto unitary synaptic connectivity evoked by single spikes or brief voltage steps (see Materials and methods): MTC activation triggered robust postsynaptic depolarization of FSIs but not RSIs, while FSI but not RSI spiking evoked inhibitory postsynaptic currents (IPSCs) in voltage-clamped MTCs and inhibitory postsynaptic potentials (IPSPs) in current-clamped MTCs (Figure 1O-R). Across 145 pairs examined, MTCs were exclusively inhibited by FSIs (Figure 1T) while 69% of FSIs exhibited postsynaptic excitation (Figure 1S) and only 1 of 27 RSIs exhibited significant excitation that was orders of magnitude weaker (Figure S6). Lack of MTC–RSI connectivity could not be attributed to any distance-dependent slicing artifact, as MTC–RSI pairs exhibited modestly shorter intersomatic distances than MTC–FSI pairs (Figure S7A). Glutamate receptor antagonists NBQX and AP5 reversibly blocked unitary MTC-to-FSI excitation, while additional application of GABA_A_ receptor (GABA_A_R) antagonist gabazine reversibly blocked unitary FSI-to-MTC inhibition (Figure S8), confirming that unitary MTC–FSI interactions are mediated by direct glutamatergic and GABAergic transmission. In total, physiological, morphological, neurochemical, and synaptic differences thus subdivide EPL-INs into two subtypes with profoundly distinct circuit functions.

The majority of MTC–FSI pairs with any connection proved reciprocally connected (Figure 1V). Unitary connectivity could further be detected between FSIs and MTCs with distal apical dendritic tufts truncated at the glomerular layer (Figure S9), confirming that MTC–FSI connectivity is localized to infraglomerular layers. No electrical coupling was detected between MTCs and FSIs (see Materials and methods). Across the EPL, unitary connection probability and strength did not differ between FSIs and mitral cells (MCs) vs. tufted cells (TCs) (Figure S10). FSIs are thus capable of influencing both streams of olfactory processing separately supported by TCs and MCs (Fukunaga et al., 2012; Igarashi et al., 2012; Geramita et al., 2016; Chae et al., 2022).

Notably, the unitary synaptic output of both MTCs and FSIs exhibited short latencies, low jitter, and high release probability (Figure S11A-C), signatures of high-fidelity, synchronous synaptic release that markedly differ from the low-fidelity, asynchronous release predicted from GCs (Ona-Jodar et al., 2020). Further of note, unitary FSI-to-MTC inhibition exhibited consistently shorter latencies than unitary MTC-to-FSI excitation (Figure S11A), potentially indicating distinct presynaptic Ca^2+^ channel expression (Hefft and Jonas, 2005). Unitary MTC-to-FSI excitation also exhibited higher release probabilities on average than unitary FSI-to-MTC inhibition (Figure S11C), though prodigious spontaneous EPSP (sEPSP) rates in FSIs (Table S1) likely masked some MTC release failures. Unitary EPSP (uEPSP) and unitary IPSC (uIPSC) amplitudes also positively correlated among reciprocally-connected MTC–FSI pairs (Figure S11D), suggesting coordinated scaling of synaptic strength.

In total, our results thus identify FSIs as a major EPL-IN subtype with strong potential to directly influence OB output through prevalent, reciprocal, and fast unitary synaptic interactions with MTCs. We therefore focused on MTC–FSI interactions throughout the remainder of the study.

### FSIs perisomatically innervate MTCs with noncanonical architecture

Post hoc inspection of reciprocally-connected MTC–FSI pairs filled with Neurobiotin (NB) revealed FSI dendritic varicosities apposed to MTC somata, proximal apical dendrites, and axon hillocks (Figure 1A,C) – subcellular domains matching the description of perisomatic innervation of projection neurons elsewhere in the brain (Freund and Katona, 2007). Post hoc multicolor confocal microscopy of an additional subset of reciprocally-connected MTC–FSI pairs filled with Lucifer Yellow (LY) and NB, respectively, likewise resolved putative contact between FSI dendrites and MTC perisomatic domains (Figure 2A,B). These patterns directly complement previous ultrastructural investigations of PV^+^ EPL-INs (Toida et al., 1994; 1996; Crespo et al., 2002), suggesting that FSIs perisomatically innervate MTCs to support similar circuit operations as perisomatic-innervating interneurons in other circuits. Two lines of evidence further suggest that FSIs *preferentially* innervate perisomatic over distal MTC domains.

**Figure 2.**
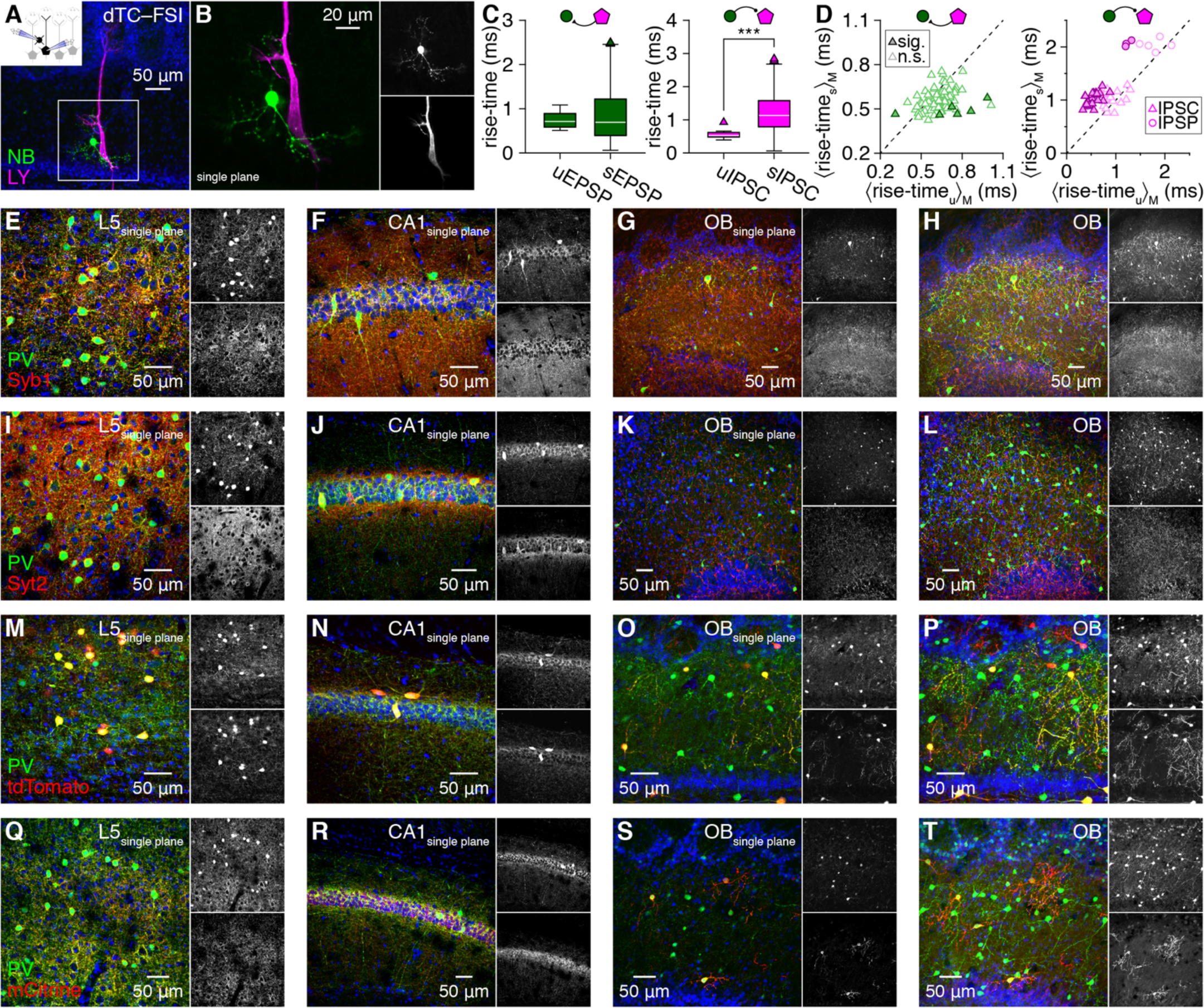
FSIs mediate fast MTC inhibition through perisomatic innervation lacking basket-like structures. **A,B:** Example reciprocally-coupled MTC–FSI pair filled with LY and NB (**A**), with single optical plane magnification (**B**). **C:** MTC-to-FSI uEPSPs (n=23 event waveforms) exhibited comparable rise-times as sEPSPs (n=613 event waveforms) in the pair in **A,B** (left; p=0.8, Wilcoxon rank-sum test with Bonferroni-corrected significance level of 0.05/78), while FSI-to-MTC uIPSCs exhibited faster rise-times than spontaneous IPSCs (sIPSCs) (right; ***p=5.3×10^−11^, Wilcoxon rank-sum test with Bonferroni-corrected significance level of 0.05/52). **D:** Comparison of unitary and spontaneous synaptic event kinetics as in **C** (only median rise-times shown for visual clarity) for 78 pairs exhibiting unitary MTC-to-FSI excitation (left) and 52 pairs exhibiting unitary FSI-to-MTC inhibition (right). **E,F:** Single optical confocal planes of layer 5 (L5) primary visual neocortex (**E**) and hippocampal CA1(**F**) revealing colocalization of Syb1 and PV in basket-like outlines of putative pyramidal cells. **G,H:** Single optical confocal plane (**G**) and maximum-intensity projection (∼50 μm depth) of Syb1 and PV in the OB. **I-L:** Same as **E-H** for Syt2 and PV. **M-T:** Same as **E-H** for PV and Cre-dependent expression of either cytosolic tdTomato (**M-P**) or membrane-localized mCitrine (**Q-T**) in PV-IRES-Cre mice.

First, connected MTC–FSI pairs exhibited significantly shorter intersomatic distances than unconnected pairs (Figure S7B-D), with connected distances within the range for FSI dendrites (radially extending ∼100 μm; Figure 1N) to contact MTC somata. This distance-dependence would be unlikely to emerge from our data if FSIs innervated distal and proximal MTC domains equally. Further, neither uEPSP nor uIPSC amplitudes correlated with distance (Figure S7E,F), suggesting that the preferential connectivity observed between nearby cells did not reflect a technical inability to resolve potentially weaker distal connections. While additional pair recordings will ultimately be required to assess to what degree FSIs may interact with the full ∼1 mm extent of MTC lateral dendrites, high rates of connectivity between nearby MTCs and FSIs together with compact FSI dendritic arbors provides a morphological foundation for preferential FSI innervation of perisomatic MTC domains.

Second, unitary FSI-to-MTC inhibitory events exhibited significantly faster rise-times than spontaneous events in 52% of connected pairs and significantly slower rise-times in 0% (Figure 2C,D), consistent with FSIs innervating MTCs at more electronically-proximal domains than inhibition from other interneurons, such as GCs (Rall, 1967). In comparison, MTC-to-FSI uEPSP rise times were significantly faster than sEPSPs in only 1% of connected pairs and significantly slower in only 6% (Figure 2C,D), suggesting that MTCs comparably innervate proximal and distal domains of FSIs. While other factors may contribute to faster MTC inhibition by FSIs than other interneurons (see Discussion), the combined observation of faster events, MTC–FSI connectivity at short intersomatic distances, and apposition of FSI dendritic varicosities to postsynaptic MTC somata, proximal apical dendrites, and axon hillocks complements prior ultrastructural findings to strongly suggest that FSIs preferentially mediate perisomatic MTC inhibition.

In neocortical and hippocampal circuits, perisomatic inhibition of pyramidal cells is mediated by profuse basket-like innervation of somata by PV^+^ axons of the local population of fast-spiking basket cells. To investigate whether the evidence of perisomatic MTC innervation by single PV^+^ FSIs presented above collectively manifests in similar canonical patterns of basket-like innervation, we immunostained for Synaptobrevin-1 (Syb1; also known as VAMP1) and Synaptotagmin-2 (Syt2), two proteins co-expressed with PV in neocortical and hippocampal basket cell axonal boutons (Sommeijer and Levelt, 2012; Vuong et al., 2018).

Consistent with prior results, Syb1 and Syt2 colocalized with PV in canonical basket-like outlines of PV^−^ putative pyramidal cells in single optical confocal planes of both neocortex and hippocampus (Figure 2E,F,I,J). In the same sections, we detected clear expression of both Syt2 and Syb1 in the OB, in contrast to reports suggesting negligible OB expression (Trimble et al., 1990; Pang et al., 2006). However, while Syb1 colocalized with PV throughout the EPL, particularly within large PV^+^ somata likely corresponding to a sparse subset of short-axon cells (Kosaka and Kosaka, 2008), no basket-like innervation of MTCs by PV/Syb1-coexpressing processes was evident in either single optical confocal planes or maximum-intensity projections (Figure 2G,H). Syt2, in turn, was also found throughout the EPL, as well as in prominent clusters throughout the mitral cell and internal plexiform layers (Figure 2K,L), potentially suggesting perisomatic MTC innervation via Chandelier-like structures (Compans and Burrone, 2023). However, these Syt2 structures not only failed to clearly associated with MTC axon initial segments (Figure S12) but further exhibited even more limited colocalization with PV than did Syb1 (Figure 2K,L).

While immunostaining for PV and presynaptic basket cell proteins Syt2 and Syb1 thus failed to reveal canonical basket-like innervation patterns, FSIs may innervate MTC somata with alternative presynaptic proteins and fine dendritic processes that stain poorly for PV. Indeed, comparison of intracellular NB and post hoc PV immunostaining revealed incomplete PV staining of fine FSI dendrites (Figure 1J; Figure S3). We therefore additionally explored whether genetic labeling of PV^+^ interneurons using crosses of PV-IRES-Cre mice and Cre-dependent cytosolic tdTomato or membrane-localized mCitrine reporter mice may reveal basket-like innervation of MTCs. In neocortex and hippocampus, cytosolic tdTomato brightly labeled PV^+^ somata and, in hippocampus, also outlined PV^−^ putative pyramidal cell somata (Figure 2M,N), while membrane-localized mCitrine could not clearly be visualized in PV^+^ somata but extensively colocalized with PV in basket-like innervation structures (Figure 2Q,R). Surprisingly, PV-IRES-Cre mice crossed to either reporter only sparsely labeled PV^+^ EPL-INs in the OB, in addition to PV^+^ periglomerular cells and deep-short axon cells (Figure 2O,P,S,T) – a level of labeling inefficiency similar to some other brain regions (Nigro et al., 2021). Nevertheless, both reporters revealed fine dendritic processes of the sparsely-labeled EPL-INs, but still failed to uncover signs of even partial basket-like MTC innervation.

Collectively, our results thus reveal kinetically-fast perisomatic inhibition of MTCs by FSIs as a prominent feature of the mouse OB, uncovering new potential for functional parallels between OB and cortical circuits. This common functional feature surprisingly emerged through noncanonical architecture void of basket-like somatic innervation, however, highlighting key structural differences between basket cells and anaxonic FSIs that may augment established computational roles of perisomatic inhibition with the potential for independent subcellular processing within release-competent FSI dendrites (see Discussion).

### FSI detonation mediates high-fidelity inhibition

Recurrent MTC inhibition powerfully regulates odorant discrimination and olfactory perception (Egger and Kuner, 2021) and is widely accepted to be slow and asynchronous. This low-fidelity signaling has been traced to both intrinsic and synaptic properties of GCs (Burton, 2017), including asynchronous GC output onto distal MTC lateral dendrites (Ona-Jodar et al., 2020) as well as weak MTC excitation of GCs, with GC spiking requiring temporal summation across prolonged MTC spike trains or the synchronous spiking of 9-30 MTCs (Pressler and Strowbridge, 2017; Mueller and Egger, 2020; Pressler and Strowbridge, 2020). In contrast to such weak GC excitation, unitary MTC-to-FSI excitation triggered FSI spiking with short latency (2.20±0.08 ms) and low jitter (0.36±0.06 ms, n=12) in 23% of connected pairs (Figure 3A,D; spike probability: 0.43±0.08 [n=19]). Identical activity was further observed spontaneously in cell-attached recordings preceding whole-cell access, with MTC spiking typical of glomerulus-wide long-lasting depolarizations (De Saint Jan et al., 2009) reliably followed by short-latency FSI spiking (Figure 3B,C), thus excluding the possibility that intracellular dialysis may have artifically elevated FSI excitability.

**Figure 3.**
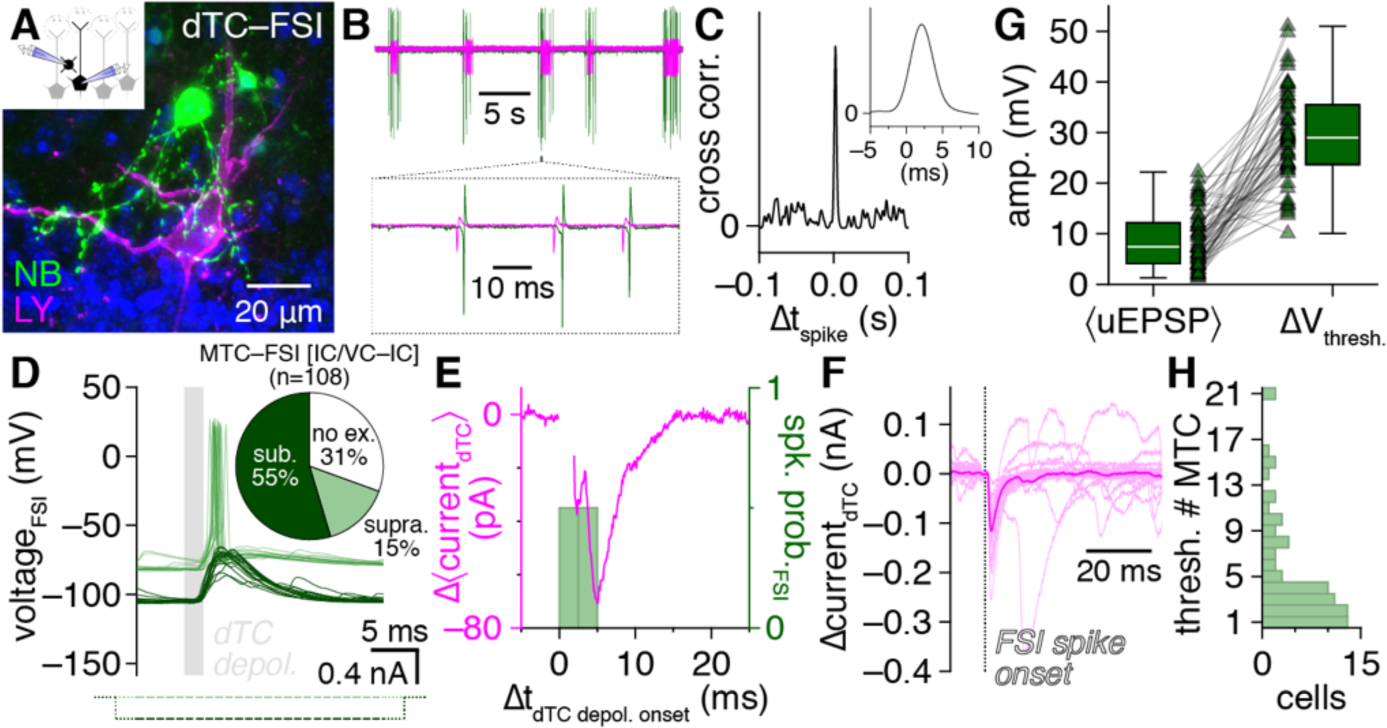
FSI detonation mediates high-fidelity recurrent MTC inhibition. **A-C :** Cell-attached recording (**B**) preceding whole-cell access of an example MTC–FSI pair (**A**) showing FSI spikes reliably following spontaneous MTC spikes by ∼2 ms (**B,C** insets). **D:** Unitary MTC release triggered FSI detonation (light green) in the pair in **A**. Negative current injection (dark green) on interleaved trials blocked detonation, revealing an underlying uEPSP. Inset: proportion of MTC–FSI pair recordings showing FSI detonation, subthreshold excitation, or no excitation. **E:** Subtraction of mean MTC currents across FSI detonation vs. uEPSP trials from **D** isolates IPSCs time-locked to FSI detonation. Voltage-step currents blanked for visual clarity. **F:** Unitary FSI-to-MTC inhibition from the pair in **A**. **G:** Comparison of mean MTC-to-FSI uEPSP amplitudes with ΔV_thresh._ across 72 FSIs with detonation probability <1. **H:** Estimated number of synchronously-spiking MTCs needed to activate each FSI (ratio of values in **G** for FSIs with detonation probability <0.5; value set to 1 for FSIs with detonation probability ≥0.5).

Such rapid unitary postsynaptic activation, or *detonation* (McNaughton and Morris, 1987), combined with the synchronous release properties of FSIs suggests that unitary MTC release under physiological conditions may trigger high-fidelity recurrent inhibition. To investigate this, we hyperpolarized FSIs in reciprocally-connected MTC–FSI pairs through negative current injection on interleaved trials, blocking detonation and revealing massive underlying uEPSPs (Figure 3D), and then examined the difference in mean MTC currents across detonation vs. uEPSP trials. Of note, each MTC can reciprocally interact with up to an estimated 10^4^ GCs (Egger and Urban, 2006; Aghvami et al., 2022), suggesting that resolving the recurrent feedback mediated by any one interneuron through somatic recording may prove exceedingly difficult unless such inhibition is particularly strong and reliable. Strikingly, our subtraction procedure indeed isolated clear recurrent IPSCs strongly timelocked to FSI detonation latencies (Figure 3E). Subtraction-isolated MTC inhibition was further comparable to the unitary FSI-to-MTC inhibition measured separately using precisely-timed current injection (Figure 3F), confirming that FSI detonation alone could fully account for the difference in mean MTC currents across detonation vs. uEPSP trials. Equivalent results were further observed when comparing spontaneously alternating detonation vs. uEPSP trials for FSIs with detonation probability <1 (Figure S13). FSI detonation is thus widespread throughout the OB and mediates strong, high-fidelity recurrent MTC inhibition.

While FSI detonation was surprisingly prevalent, many FSIs still responded to unitary MTC release with subthreshold excitation (Figure 3D). To comprehensively understand how the total population of FSIs contributes to MTC inhibition, we therefore defined ΔV_thresh._ as the difference in rheobase spike threshold and median membrane potential prior to unitary excitatory input for each FSI in our dataset exhibiting a uEPSP (Figure 3G), and then divided ΔV_thresh._ by the mean uEPSP amplitude to estimate how many synchronously-spiking MTCs are needed to activate each FSI (assuming identical unitary strength and passive EPSP summation) (Figure 3H). Mean uEPSP amplitudes exceeded ΔV_thresh._ in some FSIs (Figure 3G), consistent with a susbet of FSIs with detonation probability <1, as well as potential differences in FSI excitation by synaptic input vs. somatic current injection. Strikingly, the majority of FSIs were estimated to activate following synchronous unitary release from only 1-4 MTCs (Figure 3H), suggesting that FSIs prominently contribute to OB inhibition following even sparse synchronization of MTCs.

Similar to recurrent inhibition, lateral inhibition between heterotypic MTCs (i.e., connected to distinct glomeruli) is believed to support olfactory perception through key circuit operations, such as the binding of distinct percepts through the synchronization of MTC spiking (Kashiwadani et al., 1999). While theoretical studies specifically point to fast perisomatic lateral inhibition as a powerful synchronizing force in the OB (McTavish et al., 2012; McIntyre and Cleland, 2016) as in other circuits (Tiesinga et al., 2008; Hu et al., 2014), the few studies directly recording MTC pairs have consistently reported lateral inhibition to be slow and asynchronous (Isaacson and Strowbridge, 1998; Urban and Sakmann, 2002; Galán et al., 2006; Kapoor and Urban, 2006). As with measures of recurrent inhibition, however, these studies evoked lateral inhibition with prolonged MTC activation, often in low Mg^2+^.

Strong, high-fidelilty FSI inhibition of MTCs combined with widespread FSI detonation in physiological Mg^2+^ suggests that single MTC spikes should instead evoke detectable *fast* lateral inhibition in nearby MTCs. Consistent with this hypothesis, re-examination of MTC pair recordings from a recent datset (Burton and Urban, 2021) indeed revealed that single MTC spikes reliably evoked single, short-latency IPSPs (<10 ms) in a large proportion of nearby MTCs, including both uni- and bi-directional connections (Figure 4A-E).

**Figure 4.**
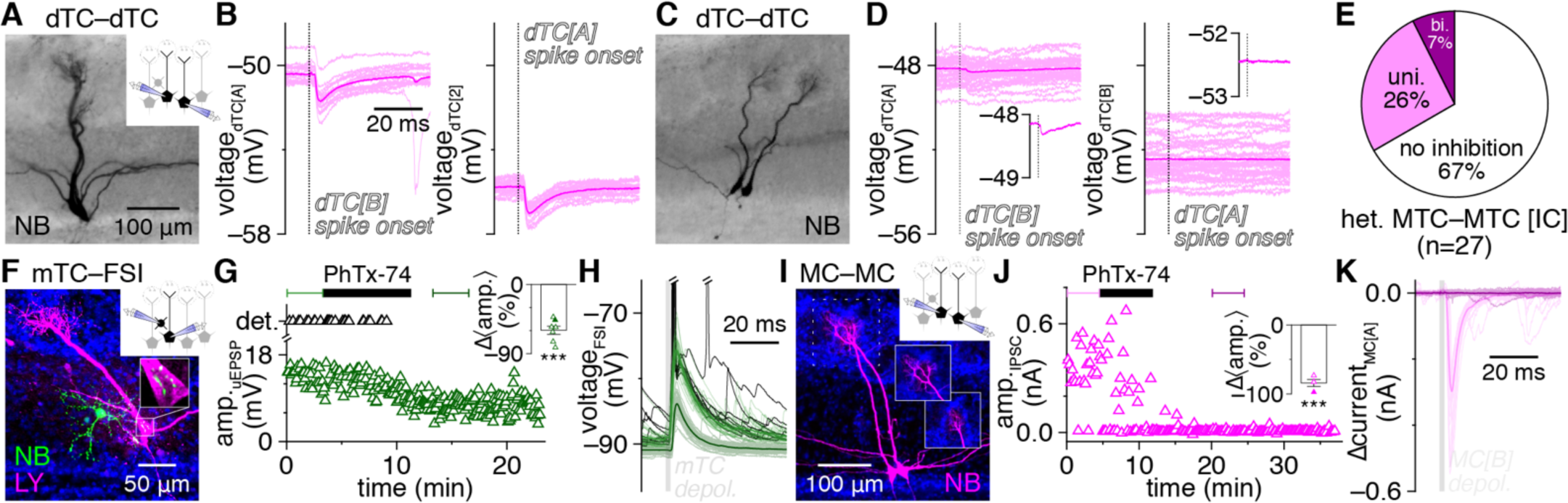
FSIs mediate fast lateral inhibition between nearby heterotypic MTCs. **A-D :** Lateral synaptic interactions evoked by single spikes within example heterotypic MTC pairs (**A**,**C**) from Burton and Urban (2021), revealing bidirectional (**B**) and unidirectional (**D**) fast lateral inhibition. Insets: magnification of mean postsynaptic voltages. Single widefield images shown. **E:** Distribution of bidirectional, unidirectional, or no fast lateral inhibitory connectivity among heterotypic MTCs recorded in current-clamp (IC). **F,G:** Example MTC–FSI pair (**F**) showing detonation occurrence and MTC-to-FSI uEPSP amplitudes before and after bath application of PhTx-74 (10 μM) (**G**). Inset: PhTx-74 significantly reduced uEPSP amplitudes (***p=3.4×10^−5^, t_6_=11.0, t-test) across 7 MTC–FSI pairs. Filled symbol corresponds to example pair. **H:** Postsynaptic FSI voltages from the pair in **F**. Light green and black traces correspond to bracketed trials (uEPSP and detonation, respectively) before PhTx-74 application in **G**; dark green traces correspond to bracketed trials after PhTx-74 application. **I,J:** Example MTC pair (**I**) showing short-latency (<10 ms) lateral IPSC amplitudes evoked by 2-ms presynaptic depolarization before and after PhTx-74 application (**J**). Inset in **I**: sub-projection of dashed box in **I** across different depths, showing apical dendrites innervating distinct glomeruli. Inset in **J**: PhTx-74 significantly reduced fast lateral IPSC amplitudes (***p=4.6×10^−4^, t_3_=16.8, t-test) across 4 connections. Filled symbol corresponds to example pair. **K:** Postsynaptic MTC currents from the pair in **I**. Dark and light traces correspond to the bracketed trials before and after PhTx-74 application in **J**, respectively.

Such fast lateral inhibition is inconsistent with the weak excitation and projected asynchronous and NMDAR-dependent output of GCs (Lage-Rupprecht et al., 2020). However, GCs strongly outnumber EPL-INs and are widely accepted to be capable of subthreshold GABA release following even weak excitation (Burton, 2017), and thus may concievably still drive fast lateral inhibition. Therefore, to provide a complementary test of the whether FSIs vs. GCs mediate fast lateral inhibition, we capitalized on the established presence vs. absence of philanthotoxin-sensitive GluA1-containing AMPARs in PV^+^ EPL-INs vs. mature GCs, respectively (Petralia and Wenthold, 1992; Giustetto et al., 1997; Petralia et al., 1997; Montague and Greer, 1999; Hamilton et al., 2008; Darcy and Isaacson, 2010; Kato et al., 2013). Philanthotoxin-7,4 (PhTx-74) indeed markedly reduced MTC-to-FSI uEPSP amplitudes, abolished FSI detonation (Figure 4F-H), and concurrently disrupted fast lateral inhibition in MTC pairs recorded in voltage-clamp to enahnce the detection of inhibition (Figure 4I-K). Our pharmacological results thus provide complementary evidence that FSI detonation plays a central role in mediating fast lateral inhibition.

The degree to which FSI-mediated fast lateral inhibition shapes OB activity, and in particular synchronization of MTC spiking, will critically depend on its prevalence relative to established forms of slow, presumably GC-mediated lateral inhibition following prolonged MTC activation. While our results so far suggest that fast lateral inhibition may be as prevalent as the ∼10% of MTC pairs exhibiting slow lateral inhhibition (Isaacson and Strowbridge, 1998), these data were collected under distinct conditions. We therefore next tested for the occurrence of fast vs. slow lateral inhibition within MTC pairs using short (2 ms) vs. long (100 ms) presynaptic voltage steps to 0 mV to trigger single spike-vs. spike train-equivalent MTC output, respectively. We defined fast lateral inhibition as a significanat increase in IPSC probability within 10 ms following presynaptic MTC activation, while slow lateral inhibition was defined as a significant increase in IPSC probability in at least three 10 ms-bins within 250 ms following presynaptic MTC activation, reflecting an asynchronous barrage of inhibition. This approach resolved clear instances of both fast and slow lateral inhibition (Figure 5A-C and Figure 5D-F, respectively), and across the population of pairs tested, fast lateral inhibition indeed proved equally as prevalent as slow lateral inhibition (Figure 5G).

**Figure 5.**
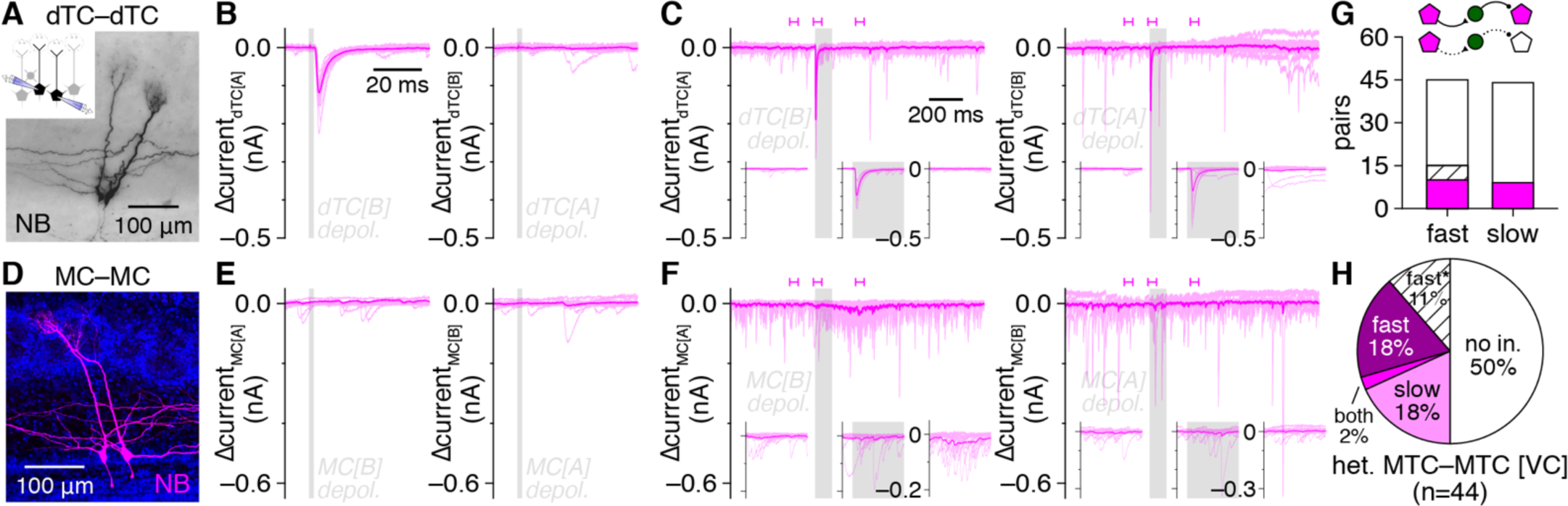
Fast lateral inhibition is a prevalent signaling mode in the OB. **A-C:** Example heterotypic MTC pair (**A**) exhibiting unidirectional fast lateral inhibition in response to 2-ms presynaptic depolarization (**B**) and bidirectional fast lateral inhibition in response to 100-ms presynaptic depolarization (**C**). Insets: magnification of postsynaptic currents corresponding to bracketed times in **C**. **D-F:** Same as **A-C** for an example heterotypic MTC pair (**D**) exhibiting unidirectional slow lateral inhibition (**F**) but no fast lateral inhibition (**E**). **G:** Heterotypic MTC pairs exhibiting fast lateral inhibition were equally as prevalent as pairs exhibiting slow lateral inhibition (p=0.84, χ^2^_[1]_=4.1×10^−2^, χ^2^ test). **H:** Distribution of fast, slow, both, or no lateral inhibition within heterotypic MTC pairs recorded in voltage clamp (VC). Pairs exhibiting fast lateral inhibition only at the start of 100-ms presynaptic depolarization (fast*) shown in hatched bars in **G,H**.

Within individual MTC pairs, fast and slow lateral inhibition appeared to occur independently. Specifically, MTC pairs exhibiting slow lateral inhibition in response to long voltage steps typically failed to exhibit fast lateral inhibition in response to short voltage steps (Figure 5D-F), while MTC pairs exhibiting fast lateral inhibition in response to short voltage steps typically continued to exhibit only fast lateral inhibition in response to long voltage steps (Figure 5A-C). Our results are thus consistent with the two lateral inhibitory signaling modes emerging through distinct circuit mechanisms: FSI-mediated fast lateral inhibition, and presumably GC-mediated slow lateral inhibition. One MTC pair exhibited both fast and slow lateral inhibition (Figure 5H), a prevalence consistent with independent occurrence of the two signaling modes (i.e., equal to the product of fast and slow lateral inhibitory prevalences) and suggesting that FSIs and GCs do not innervate distinct MTC subpopulations. Also of note, in a subset of connections (Figure 5G,H), fast lateral inhibition was only detected at the beginning of long presynaptic voltage steps (cf. Figure 5B,C right); additional experiments are needed to determine whether such signaling reflects FSI activation by rapidly summating excitatory input. In total, our results thus reveal FSI-mediated fast lateral inhibition to be highly prevalent throughout the OB in both absolute terms and relative to presumably GC-mediated slow lateral inhibition.

Odorants evoke prominent gamma-frequency oscillations throughout the OB (Kay, 2014), reflecting synchronization of MTC spiking (Kashiwadani et al., 1999; Schoppa, 2006; Burton and Urban, 2021). The fast lateral inhibition identified in our current results represents a previously-unrecognized signaling mode that may support such MTC synchronization. In turn, our estimates of MTC synchronization required to activate FSIs (Figure 3H) further suggests that synchronization of only a few MTCs should activate the majority of postsynaptic FSIs and, consequently, trigger fast lateral inhibition onto an increasing fraction of nearby MTCs to potentially spread network synchronization.

To directly test the sensitivity of FSI signaling to such MTC synchronization, we therefore next recorded MTC quartets and examined the dependence of fast lateral inhibition on the number of synchronously-activated MTCs. To limit the potential contribution of glomerular layer circuits to the recorded currents, analysis was confined to postsynaptic responses in MTCs heterotypic to all other MTCs of the quartet (as in above MTC pair recordings). In some MTCs, fast lateral inhibition occurred upon activation of one MTC regardless of whether other MTCs were coactivated (e.g., Figure 6B upper), a pattern consistent with FSI detonation and no additional FSI activation with MTC synchronization. In such instances, quantification of the mean lateral IPSC amplitude as a function of the number of synchronously-activated MTCs closely followed linear predictions (i.e., multiplying the mean singular response by the number of MTCs coactivated) (Figure 6C left). Other MTCs, however, exhibited supralinear increases in fast lateral inhibition, reflecting either greater-than-predicted increases in IPSC amplitude (e.g., Figure 6B lower, Figure 6C right) or the occurrence of fast lateral inhibition only following coactivation of 2 or 3 MTCs (e.g., Figure 6E). Across all possible connections, both the strength and prevalence of fast lateral inhibition increased supralinearly (Figure 6F,G), consistent with activation of most FSIs by synchronization of just a few MTCs. Our results thus point toward a positive feedback loop between MTC synchronization and fast lateral inhibition as a potential circuit operation supported by FSIs.

**Figure 6.**
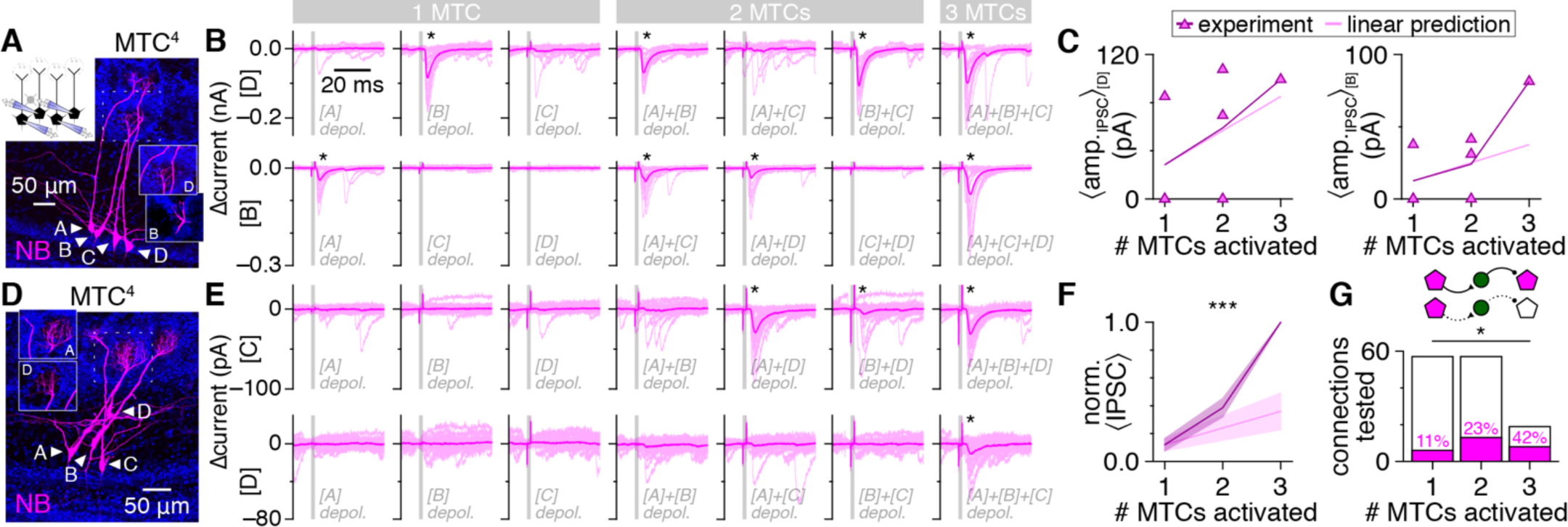
Sparse MTC synchrony supralinearly enhances fast lateral inhibition strength and prevalence. **A,B:** Example MTC quartet (**A**; inset: sub-projection of dashed box across different depths, showing MTCs [B] and [D] innervating distinct glomeruli), showing postsynaptic currents (**B**) in MTCs [D] (upper) and [B] (lower) following singular or synchronous activation of all other quartet MTCs. Asterisk: significant fast lateral inhibition. **C:** Lateral IPSC amplitudes in MTCs [D] (left) and [B] (right) from the quartet in **A** as a function of the number of synchronously-activated presynaptic MTCs. Lines: mean across all combinations of each presynaptic population size. **D,E:** Same as **A,B** for a second MTC quartet. Presynaptic voltage-step artifacts are visible in postsynaptic currents of some MTCs but terminate prior to IPSC onset. **F:** Lateral IPSC amplitudes grew supralinearly with synchronous activation of additional presynaptic MTCs (two-way ANOVA, experiment/linear prediction × # MTCs activated; ***p=9.3×10^−5^, F_1,42_=18.7, experiment vs. linear prediction; ***p=1.0×10^−8^, F_2,42_=29.4, # MTCs activated; ***p=2.4×10^−4^, F_2,42_=10.2, interaction). Values normalized within each MTC by response to synchronous activation of 3 presynaptic MTCs and averaged across 19 MTCs from 7 quartets (other MTCs were homotypic to ≥1 MTC of the quartet). **G:** Fast lateral inhibition was detected across a higher proportion of connections as more presynaptic MTCs were synchronously activated (*p=1.0×10^−2^, χ^2^_[2]_=9.2, χ^2^ test).

Understanding the net contribution of FSIs to sensory processing in the OB will further require assessing the strength of FSI-to-MTC inhibition relative to total MTC inhibition following sensory activation of the OB circuit, particularly given that each MTC receives inhibitory input from an estimated 10^4^ GCs (Egger and Urban, 2006; Aghvami et al., 2022). In our final experiment, we therefore combined MTC–FSI pair recordings with optogenetic activation of olfactory sensory neuron terminals to assess the contribution of *single* FSIs to *total* network-driven MTC lateral inhibition. Specifically, we recorded MTC inhibition evoked by photostimulation of a single lateral glomerulus while, on interleaved trials, hyperpolarizing a synaptically-coupled FSI to block its feedforward excitation-evoked spiking (Figure 7A,B). Subtraction of mean MTC currents across FSI spiking vs. EPSP trials consistently revealed robust inhibitory currents time-locked to FSI spiking (Figure 7C), regardless of whether the FSI exhibited phasic or sustained activity. Across 7 MTC– FSI pairs with suprathreshold FSI responses, activation of a single FSI remarkably accounted for 47.7±12.5% of the total inhibitory charge within 250 ms of photostimulation – a typical sniff duration (Wachowiak, 2011) (Figure 7D). Thus, at least under conditions physiologically mimicking actvation of the OB with low concentration odorants (Burton et al., 2022), high-fidelity FSI signaling accounts for a substantial fraction of total network-driven MTC lateral inhibition, suggesting that FSIs likely also play a key role in sculpting MTC tuning (Yokoi et al., 1995; Tan et al., 2010).

**Figure 7.**
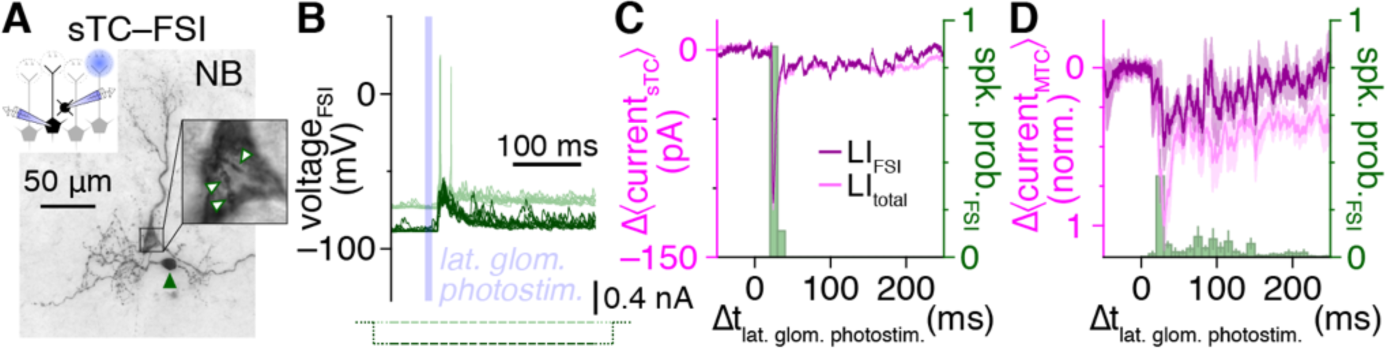
Single FSIs drive a large fraction of total network-driven lateral inhibition. **A,B:** Example MTC–FSI pair (**A**) with suprathreshold FSI response (light green) to optogenetic photostimulation of a single lateral glomerulus (**B**). Negative current injection (dark green) on interleaved trials blocked suprathreshold FSI responses. **C:** Subtraction of mean MTC currents across FSI suprathreshold vs. EPSP trials from **A** isolates lateral inhibitory currents timelocked to FSI spiking (LI_FSI_; dark magenta). Mean MTC currents across FSI suprathreshold trials, reflecting total lateral inhibition under baseline conditions (LI_total_; light magenta) shown for comparison. **D:** Subtraction-isolated lateral inhibitory currents as in **C,** normalized within each MTC by peak LI_total_ current, and averaged across MTCs of 7 MTC–FSI pairs, including 2 MTCs with distal apical dendrites truncated at the glomerular layer (Figure S9) ensuring absence of photostimulation-evoked excitatory currents.

## Discussion

To advance a foundational understanding of circuit operations in the OB supporting olfactory perception, we have systematically examined unitary synaptic interactions between MTCs and EPL-INs, a highly conserved population of anaxonic OB interneurons. Challenging the consensus that MTC inhibition is slow, low-fidelity, and primarily shaped by distributed changes in activity across a large population of lateral dendrite-innervating GCs, we found that fast-spiking EPL-INs perisomatically inhibit MTCs with release-competent dendrites and, through synaptic detonation and supralinear recruitment by sparse MTC synchronization, singularly mediate a substantial fraction of total MTC inhibition via strong, high-fidelity recurrent and lateral inhibition. These core results thus stand to reconfigure our fundamental understanding of how the OB transforms sensory input to encode olfactory information.

### EPL-IN diversity

Classic Golgi staining has established that EPL-INs are highly conserved across mammals, including at least carnivores, omnivores, and insectivores (Van Gehuchten and Martin, 1891; Schneider and Macrides, 1978; López-Mascaraque et al., 1990; Brunjes et al., 2016). Lacking functional comparison, however, these earlier studies also highlighted modest morphological differences among EPL-IN subsets of so-called Van Gehuchten, satellite, and horizontal cells. Immunohistochemistry has reinforced this apparent diversity, identifying expression of a wide variety of neurochemicals, with coexpression systematically mapped for only certain subsets (e.g., Lepousez et al., 2011; Huang et al., 2013). Collectively, these results have yielded a fairly nuanced view of EPL-INs as boutique neurons omitted from nearly all conceptual OB models.

Complementing these earlier histological approaches, we performed the first extensive physiological, morphological, neurochemical, and synaptic investigation of EPL-INs, paralleling foundational investigation of neocortical and hippocampal interneuronal diversity (Maccaferri and Lacaille, 2003; Markram et al., 2004). We surprisingly observed only two major subtypes, FSIs and RSIs, suggesting that EPL-INs are more unified in function than previously recognized. As a caveat, more subtle heterogeneity is certain to exist. Further investigation comprehensively mapping neurochemicals across all EPL-INs together with interneuron pair recordings and more extensive post hoc staining is thus poised to uncover further contributions of EPL-INs to circuit operations in the OB, in addition to potentially overcoming limitations in genetically targeting EPL-INs (e.g., Figure 2) complicating interpretation of population-level EPL-IN manipulations.

Such investigation is likely also to establish whether the greater prevalence of FSIs than RSIs in our recordings represents a true difference in cell densities or a manifestation of recording biases. In particular, to avoid superficial granule cells or deep periglomerular cells, our recordings did not sample equally across the full EPL (Table S2). Moreover, we routinely avoided targeting the largest or smallest somata for recording, as the former were likely to include at least some of the sparse EPL short-axon cells (Kosaka and Kosaka, 2008; Kosaka et al., 2008; Kosaka and Kosaka, 2010; Liberia et al., 2012; Kosaka et al., 2020) while the latter proved difficult to differentiate from resealed blebs of severed MTC dendrites. Finally, for this dataset we abandoned recordings from a small subset of interneurons with unstable resting membrane potential. This precaution enabled us to avoid EPL-INs damaged during slice preparation as well as the sparse subset of tonically-active TH^+^ short-axon cells (Figure S4) (Pignatelli et al., 2009; Liberia et al., 2012; Kosaka et al., 2020), but may have also excluded some tonically-active EPL-INs.

### Perisomatic innervation and implications of noncanonical architecture

Previous ultrastructural investigation has identified the conserved existence of perisomatic MTC innervation by certain EPL-IN subsets, including most prominently PV^+^ EPL-INs (Toida et al., 1994; 1996; Crespo et al., 2002). Lacking functional confirmation, however, such targeting has not been integrated into current conceptual OB models. Here, we measured synaptic rise-times and consistently found unitary FSI-to-MTC inhibition to be faster than spontaneous inhibition from other sources, which, together with post hoc structural analyses of synaptically-coupled pairs, compellingly argues that FSIs perisomatically inhibit MTCs.

As a caveat, differences in GABA_A_R subunit composition may alternatively underlie the observed differences in event kinetics (Eyre et al., 2012). In the OB, faster α1 subunits distribute equally throughout the EPL while slower α3 subunits concentrate in the upper EPL (Fritschy and Mohler, 1995; Pirker et al., 2000; Panzanelli et al., 2005; Lagier et al., 2007; Eyre et al., 2012), and α1 knockout slows miniature IPSC decay (though not rise-times) in MTCs (Lagier et al., 2007). At least two points suggest that the faster rise-times of FSI-mediated vs. spontaneous inhibition are not due to differential GABA_A_R subunit composition, however. First, if faster α1 subunits selectively concentrated postsynaptic to FSIs, then the noted laminar difference in subunit expression would suggest that FSIs preferentially innervate MCs instead of TCs, which we did not observe. Second, differences in subunit expression arise predominantly across cells and not across synapses within the same cell (Panzanelli et al., 2005; Lagier et al., 2007; Eyre et al., 2012). Selective localization of faster α1 subunits postsynaptic to FSIs would thus further suggest that FSIs preferentially innervate only subsets of MTCs, which was not supported by the high connectivity rates observed.

Despite such evidence for perisomatic MTC inhibition by FSIs, we failed to detect canonical basket-like innervation patterns using several complementary strategies. Absence of such profuse perisomatic innervation likely reflects two unique features of MTC–FSI signaling. First, distinct from neocortical and hippocampal pyramidal cells, both somata and dendrites of MTCs are fully excitable and release competent (Urban and Castro, 2010). Inhibition targeted exclusively to the soma would thus be less effective in controlling MTC signaling than in the case of basket innervation of pyramidal cell somata, through which all synaptic inputs must sum in order to trigger axonal output. Second, distinct from neocortical and hippocampal fast-spiking basket cells, FSIs in the OB are anaxonic and inhibit MTCs through dendritic GABA release. Distinct metabolic and trafficking requirements of axons vs. dendrites thus limit the total cable length available for FSIs to innervate MTCs, prohibiting profuse somatic wrapping. Moreover, by housing both pre- and postsynaptic machinery, FSI dendrites must distribute in a manner facilitating both GABA release onto MTCs as well as integration of glutamatergic input from the same and other MTCs. By avoiding profuse basket innervation of individual MTC somata, FSI dendrites can likely reciprocally communicate with a larger ensemble of MTCs.

While lacking canonical basket-like structure, perisomatic inhibition of MTCs by FSIs nevertheless proved surprisingly strong, hyperpolarizing MTCs by 0.46±0.45 mV from resting potentials of – 53.0±3.3 mV (n=8), comparable to unitary basket cell hyperpolarization of hippocampal and neocortical pyramidal cells by 0.45 and 1.2 mV, respectively, from resting potentials of –55 to –60 mV in adult rats (Buhl et al., 1995; Thomson et al., 1996) and far exceeding the predicted unitary GC-to-MTC IPSP amplitude of 0.01-0.03 mV (Aghvami et al., 2022). FSIs in the OB may thus achieve comparable function as cortical basket cells through strikingly distinct synaptic architecture.

Close proximity between pre- and postsynaptic machinery at reciprocal dendrodendritic and dendrosomatic synapses between FSIs and MTCs further suggests that FSIs may also support parallel subcellular processing, with excitatory input depolarizing nearby active zones to trigger local GABA release in the absence of a global spike. Such local subthreshold GABA release would not only enable FSIs to dynamically shift the balance between recurrent and lateral inhibition among MTCs, but further regulate flexible ensembles of MTCs, dramatically augmenting the computational power of each individual FSI. At least two additional factors may further augment parallel subcellular processing within FSIs. First, FSI expression of philanthotoxin-sensitive Ca^2+^-permeable AMPARs suggests that local FSI release may be dynamically enhanced by coincident Ca^2+^ influx through postsynaptic AMPARs and presynaptic Ca^2+^ channels during MTC–FSI synchronization. Second, multiple dendritic branches of each PV^+^ EPL-IN strikingly harbor clusters of Na^+^ channels that colocalize with some features of conventional axon initial segments (Kosaka and Kosaka, 2008; Kosaka et al., 2008), suggesting that FSIs may additionally be capable of generating local dendritic spikes.

### Mechanisms and implications of high-fidelity recurrent and lateral inhibition

Our results show that FSIs powerfully contribute to both recurrent and lateral MTC inhibition, motifs widely attributed solely to GCs in the OB. That contribution of single FSIs to these signaling modes was not only detectable, but constituted a large fraction of the total inhibition recorded at the soma under our experimental conditions was remarkable given that recurrent and lateral inhibition onto any MTC should conceivably also reflect the summed input from 10^4^ GCs (Egger and Urban, 2006; Aghvami et al., 2022). Our results thus suggest that OB output, specifically reflecting MTC somatic spiking, may be dominantly shaped by the comparatively small population of FSIs. A key component underlying this outsized contribution is the surprising finding that 23% of FSIs responded to unitary MTC release with suprathreshold activation. Such prevalent synaptic detonation under baseline conditions not only distinguishes FSIs from rodent neocortical and hippocampal basket cells (Molnár et al., 2008; Campanac et al., 2013), but is further uncommon throughout the entire rodent brain, typically manifesting at specialized connections crucial for high-fidelilty circuit operation (Borst et al., 1995) and learning (Eccles et al., 1966; Vandael and Jonas, 2024). Understanding how the structure, function, and plasticity of MTC–FSI synapses may relate to these more specialized detonating connections represents a key area of future investigation.

Other functional differences between FSIs and GCs beyond postsynaptic excitation further motivate reevaluation of key tenets of recurrent and lateral MTC inhibition and OB sensory processing. In particular, while GC output can be strongly gated by centrifugal cortical input (Strowbridge, 2009), FSIs readily mediate both recurrent and lateral inhibition in acute slices with disrupted corticobulbar communication. FSIs and GCs may concordantly regulate OB sensory processing in context-independent and -dependent forms, respectively, paralleling complementary intrinsic vs. extrinsic modulation of hippocampal activity by PV^+^ vs. PV^−^ basket cells (Freund and Katona, 2007). Alternatively, FSIs may also integrate various cortical and/or neuromodulatory centrifugal input to influence OB sensory processing in a context-dependent manner distinct from that of GCs. Indeed, recent investigation by Fukunaga and colleagues suggests that perisomatic inhibition, as we here show to be a prominent signaling mode in the OB mediated by FSIs, may be pivotal in reward modulation of MC sensory responses (Lindeman et al., 2024). Direct investigation of whether FSIs receive centrifugal cortical and/or neuromodulatory inputs thus stands as an important next step in evaluating contextual modulation of OB sensory processing.

Network simulations have additionally demonstrated that high-fidelity perisomatic lateral inhibition, as mediated by FSIs, is likely to promote MTC synchronization (McTavish et al., 2012; McIntyre and Cleland, 2016), consistent with mechanisms well-established in neocortical and hippocampal circuits (Tiesinga et al., 2008; Hu et al., 2014). This, together with the demonstration that sparse MTC synchronization is sufficient to activate the majority of postsynaptic FSIs, suggests that MTC– FSI interactions may play a lead role in promoting the fast-timescale MTC synchronization driving gamma-frequency OB oscillations. Further experiments are needed to evaluate this hypothesis, however, particularly given the concurrent occurrence of presumably GC-mediated slow lateral inhibition, which itself is capable of promoting synchronization of resonant gamma-frequency MTC spiking (Galán et al., 2006; Burton et al., 2012; Burton and Urban, 2021). Integration of the present unitary synaptic and physiological FSI data into advanced biophysical OB simulations (e.g., (Li and Cleland, 2017)) will provide a powerful platform for further investigating precisely when and to what degree FSIs synchronize OB output.

## Materials and methods

### Animals

All experiments were completed in compliance with the guidelines established by the Institutional Animal Care and Use Committee of Lehigh University. Optogenetic experiments (Figure 7) used gene-targeted OMP-ChR2:EYFP mice, in which the endogenous olfactory marker protein (OMP) gene is replaced with the H134R variant of channelrhodopsin-2 fused to enhanced yellow fluorescent protein (ChR2:EYFP), driving ChR2:EYFP expression in all mature olfactory sensory neurons (Smear et al., 2011). OMP-ChR2:EYFP mice were maintained on an albino C57BL/6J background and used as heterozygotes on C57BL/6J or albino C57BL/6J background to minimize olfactory sensory neuron signaling deficits, as previously described (Burton and Urban, 2015). All other electrophysiological experiments used multiple strains of mice on the C57BL/6J background and lacking genetic labeling of interneuron populations, including: M71-IRES-Cre (Li et al., 2004), M72-IRES-ChR2:EYFP (Smear et al., 2013), M72-IRES-tauCherry (Arneodo et al., 2018), and Tbet-Cre (Haddad et al., 2013), with no difference in results between strains. Immunohistochemical experiments used similar strains of mice lacking genetic labeling of interneuron populations, in addition to a subset of experiments (Figure 2M-T) using compound heterozygous crosses of gene-targeted PV-IRES-Cre mice (Hippenmeyer et al., 2005) to either RCL-tdTomato mice (Madisen et al., 2010) or RCL-ReaChR:mCitrine mice (Hooks et al., 2015) maintained on the C57BL/6J background. Mice were socially housed when possible and maintained on a 12 h light/dark cycle with ad libitum access to food and water.

### Slice preparation

Experiments were performed in acute slices prepared from P21-28 mice (n=70), consistent with the full maturation of MTC intrinsic and synaptic properties (Maher et al., 2009; Dietz et al., 2011; Yu et al., 2015) and PV^+^/CRH-Cre^+^ EPL-IN densities (Kosaka et al., 1994; Batista-Brito et al., 2008; Garcia et al., 2016). Mice of both sexes were used. For slice preparation, mice were anesthetized with isoflurane and decapitated into ice-cold oxygenated dissection solution containing the following (in mM): 125 NaCl, 25 glucose, 2.5 KCl, 25 NaHCO_3_, 1.25 NaH_2_PO_4_, 3 MgSO_4_, and 1 CaCl_2_. Brains were isolated and acute horizontal slices (310 μm thick) were prepared using a vibratome (VT1200S, Leica Biosystems). Slices recovered for 30 min in ∼37°C oxygenated Ringer’s solution that was identical to the dissection solution except with lower Mg^2+^ concentrations (1 mM MgSO_4_) and higher Ca^2+^ concentrations (2 mM CaCl_2_). Slices were then stored at room temperature until recording. Slices were prepared at approximately the same time each day relative to the animal facility light/dark cycle.

### Electrophysiology

Slices were continuously superfused with warmed oxygenated Ringer’s solution (temperature measured in bath: 30–32°C). Tissue was visualized using infrared differential interference contrast (IR-DIC) video microscopy. Recordings were targeted to the medial MOB, where the MCL reliably appears as a uniformly compact cell layer, facilitating the differentiation of cell types. MTC cell types were identified as previously (Burton and Urban, 2014; 2021). Specifically, MTCs with ≥50% of their soma displaced above the outer edge of the MCL border were classified as TCs; remaining MTCs within the MCL were classified as MCs. TCs with somata still contacting the MCL border were classified as deep TCs (dTCs); those with somata separated from the MCL and within the lower half of the EPL were classified as middle TCs (mTCs); remaining TCs with somata separated from the MCL and within the upper half of the EPL were classified as superficial TCs (sTCs). MTC cell type was assessed and classified during live IR-DIC imaging and further verified in post hoc inspection of intracellular Neurobiotin (NB; Vector Laboratories) or Lucifer Yellow CH (LY; Thermo Fisher Scientific). For MTC pair and quartet recordings, homotypic vs. heterotypic glomerular association was determined by combined assessment of: spontaneous long-lasting depolarization and/or inward current correlation (Carlson et al., 2000), comparatively slow lateral excitation among homotypic MCs (Urban and Sakmann, 2002; Burton and Urban, 2021), and post hoc inspection of intracellular NB.

Current-clamp data were recorded using electrodes filled with (in mM): 135 K-gluconate 1.8 KCl, 8.8 HEPES, 10 Na-phosphocreatine, 4 Mg-ATP, 0.3 Na-GTP, and 0.2 EGTA. Voltage-clamp data were recorded using electrodes filled with (in mM): 131 CsCl, 3.8 K-gluconate, 0.05 KCl, 8.8 HEPES, 10 Na-phosphocreatine, 4 Mg-ATP, and 0.3 Na-GTP. Electrode solutions additionally contained 8.8 mM GABA or glutamate (for EPL-IN or MTC recordings, respectively) to preclude rundown of dendritic release (Smith and Jahr, 2002; Ma and Lowe, 2007; De Saint Jan et al., 2009; Najac et al., 2015; Zak et al., 2015), as well as either 0.025 mM Alexa Fluor 488 or 594 hydrazide to permit live visualization and 0.2% NB to permit post hoc inspection. For a subset of recordings, voltage-clamp data were recorded using electrodes filled with (in mM): 66 CsCl, 61 Cs-gluconate, 3.8 K-gluconate, 0.9 KCl, 8.8 HEPES, 8.8 glutamate, 10 Na-phosphocreatine, 4 Mg-ATP, 0.3 Na-GTP, and 0.1% Lucifer Yellow CH; these recordings were excluded from other voltage-clamp data for comparisons of absolute IPSC amplitudes across cells (Figure S7F; Figure S10E; Figure S11D). Electrode resistances were 3-10 MΩ. Current-clamped cells were held at their resting membrane potential (i.e., 0 pA holding current); voltage-clamped cells were held at –70 mV. In current-clamp recordings, pipette capacitance was neutralized and series resistance (EPL-IN: 37.0±0.9 MΩ [n=145]; MTC: 18.8±0.9 MΩ [n=54]) was compensated using the MultiClamp Bridge Balance operation. In voltage-clamp recordings, series resistance (MTC: 20.5±0.5 MΩ [n=235]) was not compensated but was monitored continuously to ensure adequate electrode access and recording quality. Current-clamp recordings of cells with unstable and/or depolarized resting membrane potential (>–50 mV) were abandoned to exclude damaged or otherwise unhealthy cells from our dataset. Recordings of MTCs with truncated apical dendrites were abandoned, except where noted. Cell-attached data were recorded prior to obtaining whole-cell access, using the same electrodes as used for subsequent current- and voltage-clamp recordings. Data were low-pass filtered at 4 kHz and digitized at 10 kHz using MultiClamp 700A and 700B amplifiers and an ITC-18 acquisition board controlled by custom software written in IGOR Pro.

To probe unitary synaptic output of EPL-INs onto MTCs, we monitored MTC voltage (n=54 pairs) or current (n=91 pairs) while injecting 1-ms suprathreshold current pulses (1.5-2.5 nA) into the EPL-IN to trigger single spikes. Voltage-clamped MTCs were recorded with a high-Cl^−^ internal solution to reverse and amplify the GABAergic driving force, enabling inhibitory postsynaptic currents (IPSCs) to be detected at hyperpolarized potentials. In the same pairs, reciprocal unitary synaptic output of the MTC onto the EPL-IN was investigated by monitoring EPL-IN voltage while injecting a 1-ms suprathreshold current pulse (1.5-2.5 nA) or a 2-ms voltage step to 0 mV (approximating the depolarization of a single spike) in the current-or voltage-clamped MTCs, respectively. Synaptic pharmacology was assessed using bath application of DL-AP5 (50 μM; Tocris, 3693), NBQX (10 μM; Tocris, 1262), Gabazine (10 μM; Tocris, 1262) and Philanthotoxin-7,4 (PhTx-74; 10 μM; Alomone Labs, P-120).

To evaluate whether MTCs and FSIs were also linked by electrical coupling, we injected hyperpolarizing step currents sequentially into each cell of a subset of synaptically-coupled MTC– FSI pairs recorded in current-clamp mode and calculated coupling coefficients. MTC step current injection (–407±64 pA) strongly hyperpolarized MTC membrane potentials (ΔV=–32.9±2.0 mV) but caused no significant change in FSI membrane potentials (ΔV=–0.02±0.08 mV; p=0.8, t_6_=–0.3, n=7, t-test), yielding an MTC-to-FSI coupling coefficient not significantly different from zero (0.13±0.26%, p=0.7, t_6_=0.5, n=7, t-test). Similarly, FSI step current injection (–220±58 pA) strongly hyperpolarized FSI membrane potentials (ΔV=–19.3±1.4 mV) but caused no significant change in MTC membrane potentials (ΔV=–0.03±0.09 mV; p=0.7, t_4_=–0.4, n=5, t-test), yielding an FSI-to-MTC coupling coefficient also not significantly different from zero (0.18±0.47%, p=0.7, t_4_=0.4, n=5, t-test).

For MTC–FSI pair recordings combined with optogenetic photostimulation (Figure 7), slices were illuminated by a 75 W xenon arc lamp passed through a YFP filter set and 60× water-immersion objective centered on a nearby lateral glomerulus (∼2-3 glomeruli away from the recorded MTC glomerulus), with field-stops closed to achieve single glomerulus activation, as previously performed (Burton and Urban, 2015). Photostimulation consisted of a 10-ms light pulse.

### Histology

For fluorescent NB labeling with post hoc immunohistochemistry, acute slices containing NB-filled cells were fixed with paraformaldehyde (PFA; 4%) in phosphate buffer (PB; 0.1 M) for >24 h at 4C, washed, and then incubated in blocking solution (2% normal serum and 0.4% Triton X-100, in PB). Slices were then incubated in blocking solution containing Alexa Fluor 488 or 594 Streptavidin (1 μg/mL) and primary antibodies, washed, incubated in blocking solution containing secondary antibodies and Hoechst 33342 (0.25 μg/ml), washed, and mounted with Fluoromount G (SouthernBiotech). For fluorescent NB labeling without post hoc immunohistochemistry, antibodies were omitted. Non-fluorescent chromogenic NB labeling was performed as previously described (Burton et al., 2017).

For immunohistochemistry, adult mice were anesthetized with intraperitoneal injection of ketamine (200 mg/kg) and xylazine (20 mg/kg) and then transcardially perfused with 0.1 M phosphate-buffered saline followed by PFA (4%) in PB. Brains were extracted, postfixed overnight, and then sectioned with a vibratome (Leica, VT1000S). Free-floating 50-μm sagittal sections were then incubated in blocking solution (2% normal serum and 0.1% Triton X-100, in PB), washed, and incubated in antibody solution (2% normal serum and 0.05% Tween 20, in PB) containing combinations of the following primary antibodies: guinea pig anti-parvalbumin (1:2,000; Synaptic Systems, 195 004), mouse anti-synaptotagmin-2 (1:200; Zebrafish International Resource Center, znp-1), rabbit anti-TRIM-46 (1:1,000; Synaptic Systems, 377 008), rabbit anti-Lucifer Yellow (1:1,000; Thermo Fisher Scientific, A-5750), rabbit anti-parvalbumin (1:2,000; Synaptic Systems, 195 002; data not shown), rabbit anti-synaptobrevin-1 (1:500; Synaptic Systems, 104 002), rabbit anti-vasoactive intestinal peptide (1:4,000; ImmunoStar, 20077), and sheep anti-tyrosine hydroxylase (1:1,000; Millipore, AB1542). Sections were then washed, incubated in antibody solution with fluorescent secondary antibodies and Hoechst 33342, washed, and mounted with Fluoromount G. All washes were performed with PB. All incubation steps took place for 1-3 h at room temperature or overnight at 4C. Secondary antibodies were used at 1:600. At least two sections from each of two male and two female mice were examined for each experiment; no gross sex-dependent differences were observed.

Brightfield and fluorescent widefield images were collected using an upright Nikon Eclipse E1000 microscope using 20× air and 100× oil-immersion objectives. Fluorescent confocal z-stacks were collected using an inverted Zeiss LSM 880 confocal microscope using a 20× air objective and 25× and 63× oil-immersion objectives, 0.5-μm z-steps, and 2048×2048 resolution. Morphological images included in figures show maximum-or minimum-intensity projections of fluorescent confocal z-stacks or chromogenic widefield z-stacks, respectively, except where noted. Neuron morphologies were reconstructed from confocal z-stacks and analyzed using the SNT plugin in ImageJ (Fiji) (Arshadi et al., 2021). Intersomatic distances were measured from single widefield fluorescent images of intracellular NB and LY, and thus do not account for potential differences in cell depth in the tissue.

### Data analysis

Values reported are either mean ± SEM or median (first quartile [Q_1_], third quartile [Q_3_]) for normally or non-normally distributed data, respectively. Line plots with thin and thick lines denote individual trials and mean, respectively. Line plots with shading denote mean ± SEM. Single, double, and triple asterisks in figures and tables denote statistical significance at p<0.05, p<0.01, and p<0.001 levels, respectively. For each statistical test, data normality was first determined by the Shapiro– Wilk test, and nonparametric tests applied where appropriate. For visual comparison of normally distributed data, all individual data points are displayed in addition to sample mean and SEM. For visual comparison of non-normally distributed data, data are displayed as standard boxplots, with data points denoting sample outliers. Significance thresholds were corrected for multiple comparisons where indicated. All recorded traces, spikes, and potential unitary synaptic events were visually inspected for detection accuracy. No differences in synaptic connectivity or intrinsic biophysical properties were detected between sex, and data were therefore pooled across male and female mice.

EPL-INs were classified as FSIs or RSIs using step current-evoked spiking responses, as shown in Figure 1B,D,F,H and reinforced by unbiased agglomerative hierarchical clustering (Figure 1I) and principal component analysis (Figure S1). Hierarchical clustering was performed on z-scored intrinsic biophysical properties using Ward’s method, with properties ordered as in Figure S1, beginning with AHP 50% decay. Given the binary connectivity profile observed among MTCs and FSIs vs. RSIs, for analysis of unitary synaptic properties the presence of a unitary connection was used as a complementary criterion to identify an additional 10 FSIs from pair recordings for which step current-evoked spiking responses were not obtained. Results did not differ if these 10 FSIs were omitted.

Resting membrane potential was recorded immediately after obtaining whole-cell access, and for EPL-INs was defined as the 10^th^ percentile of voltage recorded for each cell to limit the contribution of prodigious sEPSP rates (e.g., Figure S2A,F,K,P). Membrane time constant, input resistance, and capacitance were calculated using the voltage trajectory and maximum voltage change evoked by a 50-pA, 100-ms hyperpolarizing step current injection (e.g., Figure S2B,G,L,Q). Spontaneous firing rates were defined as the spike-count firing rate calculated from the total number of spikes not driven by evoked unitary synaptic input recorded over 151.6 s (103.2, 184.0) (n=145) for each cell.

Firing rate-current (FI) curves were examined using 500-ms depolarizing step current injections ranging from 50 to 600 pA in steps of 50 pA for FSIs and less excitable RSIs (e.g., Figure S2D,I,S), while RSIs that readily underwent depolarization block were examined using step currents ranging from 10 to 100 pA in steps of 10 pA (e.g., Figure S2N). FI curve spike times were detected using a voltage derivative threshold of 15 mV/ms. FI curve rates were calculated from the median inverse interspike interval (ISI) evoked by each step current. Spike properties (with the exception of afterdepolarization [ADP] amplitudes) were calculated from the first spike evoked by the weakest suprathreshold step current (i.e., rheobase). Spike amplitude was calculated as the difference between spike threshold (i.e., the voltage at spike onset) and spike peak. Maximum and minimum spike slopes were calculated as the maximum and minimum voltage derivatives. Spike width was calculated as the full-width at half-maximum spike amplitude. Spike afterhyperpolarization (AHP) amplitude was calculated as the difference in spike threshold and the minimum voltage reached within 10 ms of spike onset. AHP 50% decay was calculated as the latency from AHP onset (i.e., spike falling phase matching spike threshold) to decay of the AHP to 50% of its amplitude. Maximum FI curve gain was calculated as the maximum FI curve derivative. Maximum FI curve rate was calculated as the inverse of the minimum ISI detected. Relative and absolute spiking adaptation were calculated from the response to the weakest suprathreshold step current evoking sustained activity (≥5 spikes), and from the first evoked spike cluster within that response for FSIs exhibiting clustered spiking (e.g., Figure 1B,D).

Spike times evoked by 1-ms suprathreshold current pulses (1.5-2.5 nA) or unitary MTC release excitation were calculated as the time at which membrane potentials (upsampled 100-fold) exceeded –30 mV. EPL-IN spike waveforms evoked by such current pulses typically lacked an AHP and instead displayed an ADP (e.g., Figure S2C). ADP amplitudes were calculated as the difference between the minimum and maximum post-spike voltage (occurring within 6 ms of spike falling phases) for spike waveforms lacking an AHP.

Postsynaptic events were detected using a standard template-matching function in Axograph (Clements and Bekkers, 1997) with double-exponential template (Table S3). For analysis of unitary MTC–EPL-IN synaptic connectivity, presynaptic activation (either pulse-evoked spikes or brief voltage-steps) was triggered every 8-22 s and peristimulus time histograms of postsynaptic events were calculated across all trials. Unitary connectivity was classified as significant if the probability of a postsynaptic event within the first time bin following presynaptic spike time or voltage step onset exceeded the mean + the standard deviation × multiplication factor of the event probability across the 1 s preceding presynaptic spike time or voltage step onset. For tests of unitary EPL-IN excitation, 5 ms time bins were used (accommodating longer latencies following presynaptic voltage steps – see below) and the significance multiplication factor was set to 2.75; suprathreshold postsynaptic responses occurring within 5 ms of presynaptic activation were additionally used to classify a unitary connection as significant. For tests of unitary MTC inhibition, 2.5 ms time bins were used and the significance multiplication factor was set to 3. Unitary postsynaptic event latency was calculated as the latency from presynaptic spike onset to the time at which the postsynaptic response (upsampled 20-fold) reached 5% of its amplitude. Unitary FSI excitation latencies were significantly longer (2.36±0.06 ms vs. 1.17±0.04 ms; p=6.6×10^−25^, t_75_=15.4, t-test) and had higher jitter (0.47 [0.35, 0.62] vs. 0.24 [0.12, 0.26]; p=2.7×10^−7^, r.s.=2290, Wilcoxon rank-sum test) when measured from MTC depolarization onset in voltage-clamp mode (n=46) than from MTC spike onset in current-clamp mode (n=31), reflecting lower precision in identifying MTC activation timing in voltage-clamp mode. All unitary FSI excitation latencies reported in the main text, including uEPSP and detonation latencies, are thus restricted to pairs recorded in current-clamp mode with precisely-measured presynaptic spike onsets. Postsynaptic rise times were calculated as the duration for the postsynaptic response (upsampled 20-fold) to increase from 20% to 80% of its amplitude. sEPSP rates were calculated for each EPL-IN from the total number of EPSPs occurring in the 1 s preceding presynaptic activation across all trials, encompassing 39.9±1.3 s (n=145) total recording for each cell. Median sEPSP half-widths were calculated as the full-width at half-maximum amplitude of the median sEPSP waveform.

For analysis of cell-attached spontaneous spike-time synchrony (Figure 3C), spike times were detected using a current threshold of 18 pA and then convolved with a Gaussian kernel (1 ms standard deviation) to account for slight differences in spike waveforms. Trains of convolved spike times were then mean-subtracted and the cross-correlogram calculated.

For analysis of lateral inhibition among MTCs, peristimulus time histogram analysis was used as above to classify lateral inhibition as significant, with 10 ms time bins (accommodating disynaptic latencies) and a significance multiplication factor of 2 for current-clamped MTCs and 3 for voltage-clamped MTCs.

## Supplementary tables

**Table S1.**
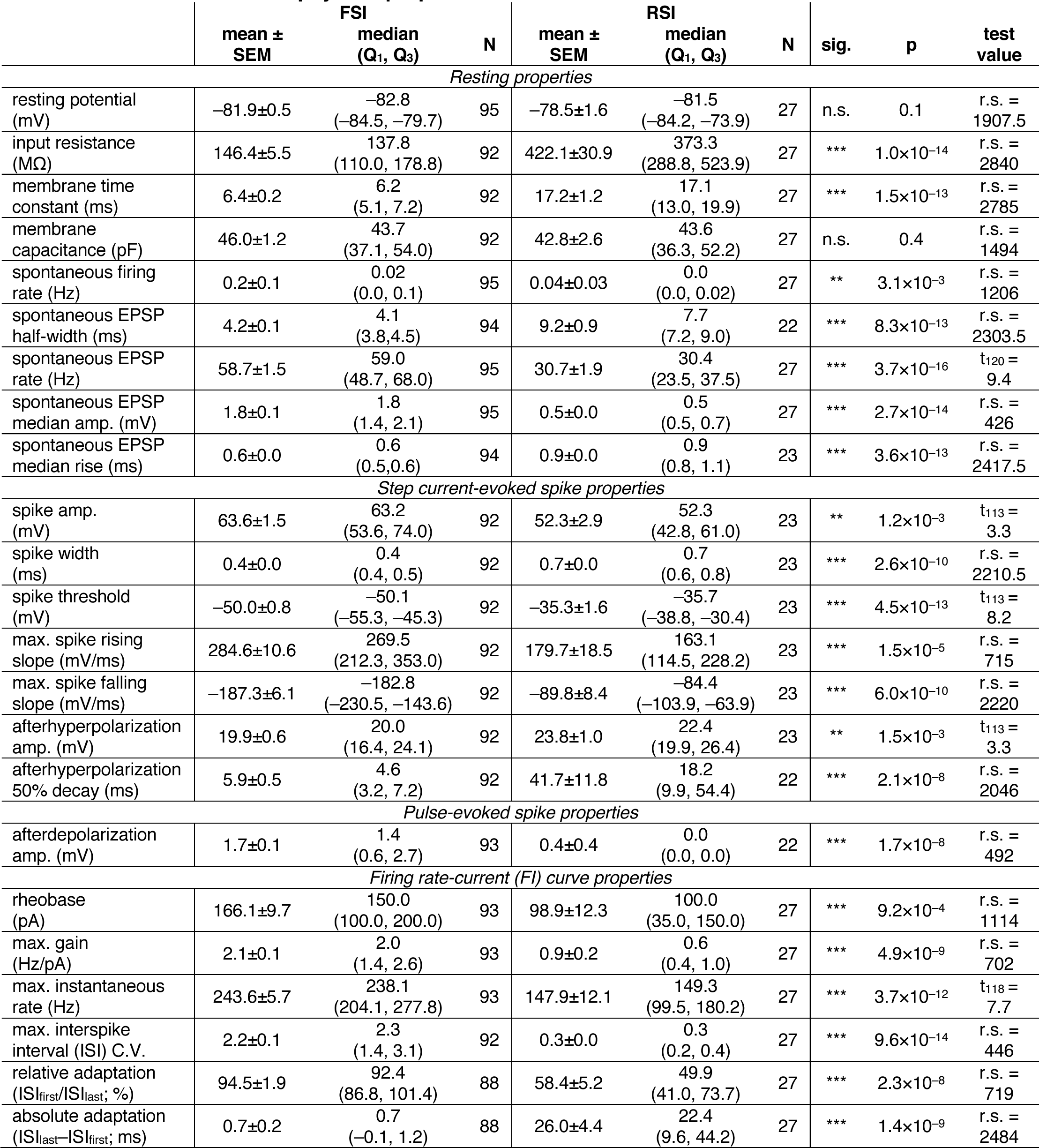
EPL-IN intrinsic biophysical properties.

|  | mean ± SEM | FSI<br>median<br>(Q<sub>1</sub>, Q<sub>3</sub>) | N | mean ± SEM | RSI<br>median<br>(Q<sub>1</sub>, Q<sub>3</sub>) | N | sig. | p | test value |
| --- | --- | --- | --- | --- | --- | --- | --- | --- | --- |
| Resting properties |  |  |  |  |  |  |  |  |  |
| resting potential (mV) | -81.9±0.5 | -82.8<br>(-84.5, -79.7) | 95 | -78.5±1.6 | -81.5<br>(-84.2, -73.9) | 27 | n.s. | 0.1 | r.s. = 1907.5 |
| input resistance (MΩ) | 146.4±5.5 | 137.8<br>(110.0, 178.8) | 92 | 422.1±30.9 | 373.3<br>(288.8, 523.9) | 27 | *** | 1.0×10<sup>-14</sup> | r.s. = 2840 |
| membrane time constant (ms) | 6.4±0.2 | 6.2<br>(5.1, 7.2) | 92 | 17.2±1.2 | 17.1<br>(13.0, 19.9) | 27 | *** | 1.5×10<sup>-13</sup> | r.s. = 2785 |
| membrane capacitance (pF) | 46.0±1.2 | 43.7<br>(37.1, 54.0) | 92 | 42.8±2.6 | 43.6<br>(36.3, 52.2) | 27 | n.s. | 0.4 | r.s. = 1494 |
| spontaneous firing rate (Hz) | 0.2±0.1 | 0.02<br>(0.0, 0.1) | 95 | 0.04±0.03 | 0.0<br>(0.0, 0.02) | 27 | ** | 3.1×10<sup>-3</sup> | r.s. = 1206 |
| spontaneous EPSP half-width (ms) | 4.2±0.1 | 4.1<br>(3.8, 4.5) | 94 | 9.2±0.9 | 7.7<br>(7.2, 9.0) | 22 | *** | 8.3×10<sup>-13</sup> | r.s. = 2303.5 |
| spontaneous EPSP rate (Hz) | 58.7±1.5 | 59.0<br>(48.7, 68.0) | 95 | 30.7±1.9 | 30.4<br>(23.5, 37.5) | 27 | *** | 3.7×10<sup>-16</sup> | t<sub>120</sub> = 9.4 |
| spontaneous EPSP median amp. (mV) | 1.8±0.1 | 1.8<br>(1.4, 2.1) | 95 | 0.5±0.0 | 0.5<br>(0.5, 0.7) | 27 | *** | 2.7×10<sup>-14</sup> | r.s. = 426 |
| spontaneous EPSP median rise (ms) | 0.6±0.0 | 0.6<br>(0.5, 0.6) | 94 | 0.9±0.0 | 0.9<br>(0.8, 1.1) | 23 | *** | 3.6×10<sup>-13</sup> | r.s. = 2417.5 |
| Step current-evoked spike properties |  |  |  |  |  |  |  |  |  |
| spike amp. (mV) | 63.6±1.5 | 63.2<br>(53.6, 74.0) | 92 | 52.3±2.9 | 52.3<br>(42.8, 61.0) | 23 | ** | 1.2×10<sup>-3</sup> | t<sub>113</sub> = 3.3 |
| spike width (ms) | 0.4±0.0 | 0.4<br>(0.4, 0.5) | 92 | 0.7±0.0 | 0.7<br>(0.6, 0.8) | 23 | *** | 2.6×10<sup>-10</sup> | r.s. = 2210.5 |
| spike threshold (mV) | -50.0±0.8 | -50.1<br>(-55.3, -45.3) | 92 | -35.3±1.6 | -35.7<br>(-38.8, -30.4) | 23 | *** | 4.5×10<sup>-13</sup> | t<sub>113</sub> = 8.2 |
| max. spike rising slope (mV/ms) | 284.6±10.6 | 269.5<br>(212.3, 353.0) | 92 | 179.7±18.5 | 163.1<br>(114.5, 228.2) | 23 | *** | 1.5×10<sup>-5</sup> | r.s. = 715 |
| max. spike falling slope (mV/ms) | -187.3±6.1 | -182.8<br>(-230.5, -143.6) | 92 | -89.8±8.4 | -84.4<br>(-103.9, -63.9) | 23 | *** | 6.0×10<sup>-10</sup> | r.s. = 2220 |
| afterhyperpolarization amp. (mV) | 19.9±0.6 | 20.0<br>(16.4, 24.1) | 92 | 23.8±1.0 | 22.4<br>(19.9, 26.4) | 23 | ** | 1.5×10<sup>-3</sup> | t<sub>113</sub> = 3.3 |
| afterhyperpolarization 50% decay (ms) | 5.9±0.5 | 4.6<br>(3.2, 7.2) | 92 | 41.7±11.8 | 18.2<br>(9.9, 54.4) | 22 | *** | 2.1×10<sup>-8</sup> | r.s. = 2046 |
| Pulse-evoked spike properties |  |  |  |  |  |  |  |  |  |
| afterdepolarization amp. (mV) | 1.7±0.1 | 1.4<br>(0.6, 2.7) | 93 | 0.4±0.4 | 0.0<br>(0.0, 0.0) | 22 | *** | 1.7×10<sup>-8</sup> | r.s. = 492 |
| Firing rate-current (FI) curve properties |  |  |  |  |  |  |  |  |  |
| rheobase (pA) | 166.1±9.7 | 150.0<br>(100.0, 200.0) | 93 | 98.9±12.3 | 100.0<br>(35.0, 150.0) | 27 | *** | 9.2×10<sup>-4</sup> | r.s. = 1114 |
| max. gain (Hz/pA) | 2.1±0.1 | 2.0<br>(1.4, 2.6) | 93 | 0.9±0.2 | 0.6<br>(0.4, 1.0) | 27 | *** | 4.9×10<sup>-9</sup> | r.s. = 702 |
| max. instantaneous rate (Hz) | 243.6±5.7 | 238.1<br>(204.1, 277.8) | 93 | 147.9±12.1 | 149.3<br>(99.5, 180.2) | 27 | *** | 3.7×10<sup>-12</sup> | t<sub>118</sub> = 7.7 |
| max. interspike interval (ISI) C.V. | 2.2±0.1 | 2.3<br>(1.4, 3.1) | 92 | 0.3±0.0 | 0.3<br>(0.2, 0.4) | 27 | *** | 9.6×10<sup>-14</sup> | r.s. = 446 |
| relative adaptation (ISI<sub>first</sub>/ISI<sub>last</sub>; %) | 94.5±1.9 | 92.4<br>(86.8, 101.4) | 88 | 58.4±5.2 | 49.9<br>(41.0, 73.7) | 27 | *** | 2.3×10<sup>-8</sup> | r.s. = 719 |
| absolute adaptation (ISI<sub>last</sub>-ISI<sub>first</sub>; ms) | 0.7±0.2 | 0.7<br>(-0.1, 1.2) | 88 | 26.0±4.4 | 22.4<br>(9.6, 44.2) | 27 | *** | 1.4×10<sup>-9</sup> | r.s. = 2484 |

**Table S2.** EPL-IN anatomical and morphometric properties.

|  | mean ±<br>SEM | FSI<br>median<br>(Q<sub>1</sub>, Q<sub>3</sub>) | N | mean ±<br>SEM | RSI<br>median<br>(Q<sub>1</sub>, Q<sub>3</sub>) | N | sig. | p | test<br>value |
| --- | --- | --- | --- | --- | --- | --- | --- | --- | --- |
| relative EPL depth<br>(0=MCL, 1=GL) | 0.59±0.02 | 0.62<br>(0.49, 0.71) | 104 | 0.65±0.02 | 0.63<br>(0.55, 0.76) | 28 | n.s. | 0.2 | r.s. =<br>6659 |
| soma area<br>(µm<sup>2</sup>) | 94.2±5.2 | 97.0<br>(77.9, 104.8) | 16 | 78.6±5.9 | 73.8<br>(61.6, 96.4) | 9 | n.s. | 0.07 | t<sub>23</sub> =<br>1.9 |
| soma max. diameter<br>(µm) | 13.2±0.4 | 13.2<br>(12.4, 13.9) | 16 | 13.3±0.8 | 12.6<br>(11.4, 15.0) | 9 | n.s. | 0.9 | t<sub>23</sub> =<br>0.1 |
| soma max. diameter/<br>min. diameter | 1.37±0.06 | 1.32<br>(1.16, 1.53) | 16 | 1.48±0.11 | 1.38<br>(1.28, 1.50) | 9 | n.s. | 0.3 | r.s. =<br>189 |
| dendritic fractal<br>dimension | 1.11±0.01 | 1.12<br>(1.08, 1.14) | 16 | 1.08±0.01 | 1.07<br>(1.05, 1.09) | 9 | * | 0.01 | t<sub>23</sub> =<br>2.8 |
| total dendritic length<br>(mm) | 1.44±0.07 | 1.48<br>(1.21, 1.68) | 16 | 0.84±0.14 | 0.71<br>(0.53, 1.14) | 9 | *** | 3.6×10<sup>-4</sup> | t<sub>23</sub> =<br>4.2 |
| # primary dendritic<br>branches | 4.3±0.5 | 4.0<br>(3.0, 5.5) | 16 | 3.9±0.6 | 4.0<br>(3.0, 4.0) | 9 | n.s. | 0.7 | r.s. =<br>214.5 |
| # terminal dendritic<br>branches | 50.8±4.0 | 49.5<br>(38.5, 63.5) | 16 | 15.7±3.8 | 11.0<br>(9.8, 21.5) | 9 | *** | 2.1×10<sup>-4</sup> | r.s. =<br>274 |
| # total dendritic<br>branches | 96.4±7.7 | 95.5<br>(72.5, 115.5) | 16 | 27.0±7.3 | 18.0<br>(15.8, 35.8) | 9 | *** | 2.0×10<sup>-4</sup> | r.s. =<br>274 |
| dendritic spine<br>density (#/100 µm) | 3.2±0.2 | 3.2<br>(2.7, 3.8) | 16 | 11.6±2.2 | 13.7<br>(4.4, 16.0) | 9 | *** | 3.1×10<sup>-5</sup> | t<sub>23</sub> =<br>5.2 |

**Table S3.** Template parameters for postsynaptic event detection.

| Event | baseline<br>(ms) | duration<br>(ms) | amplitude<br>(a.u.) | rise<br>(ms) | decay<br>(ms) | min. separation<br>(ms) | threshold<br>(×S.D.) |
| --- | --- | --- | --- | --- | --- | --- | --- |
| MTC IPSC | 0.5 | 5.0 | -1 | 0.8 | 4.0 | 0.5 | 2.5 |
| MTC IPSP | 1.0 | 8.0 | -1 | 1.0 | 6.0 | 0.5 | 3.0 |
| EPL-IN EPSP | 0.5 | 5.0 | 1 | 0.7 | 4.0 | 0.5 | 2.5 |

## Supplementary figures

**Figure S1.**
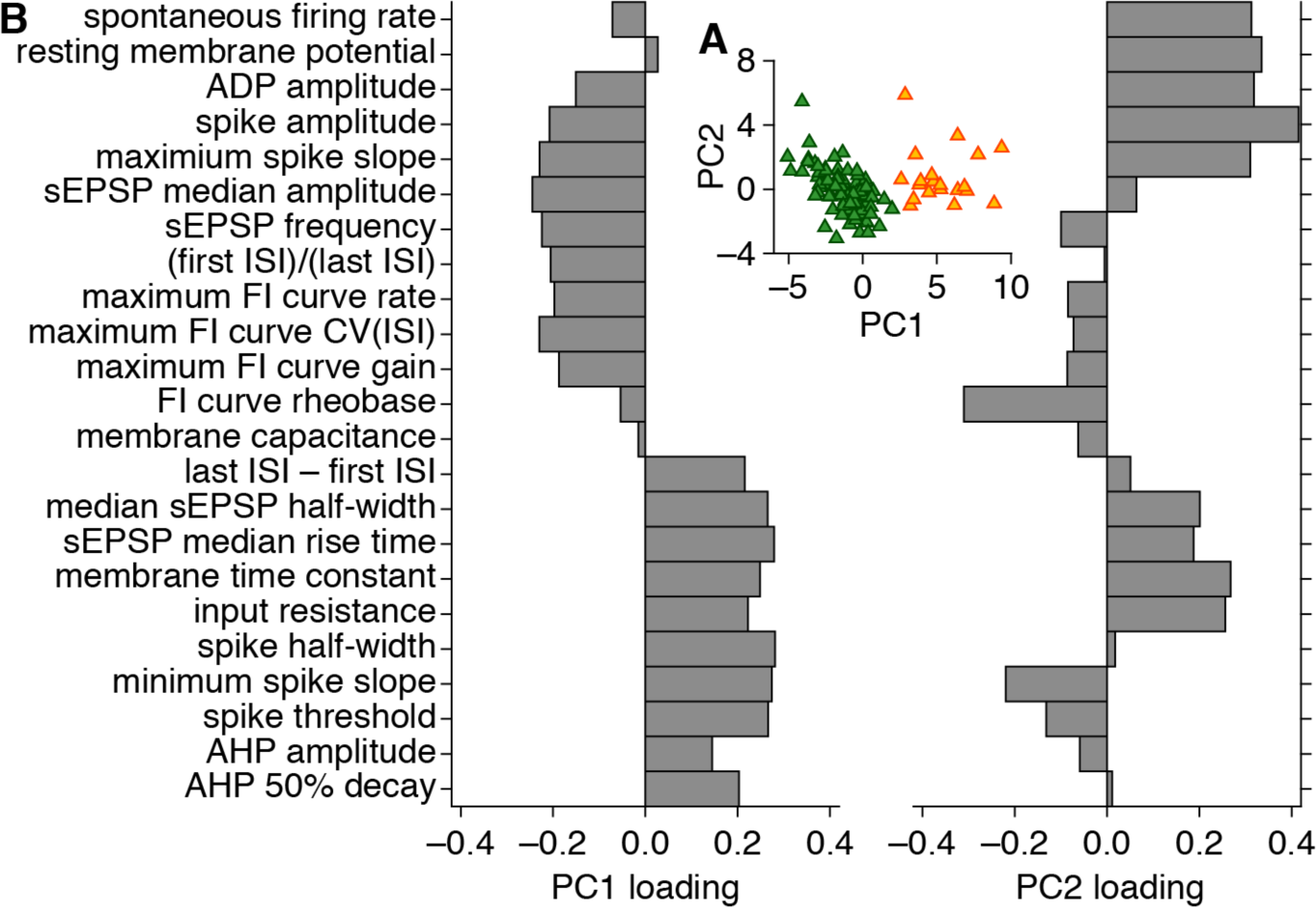
Principal component analysis reinforces subdivision of EPL-INs into FSIs and RSIs. **A:** Projection of EPL-INs onto the first two principal components (PC1 and PC2) defined by principal component analysis of z-scored intrinsic biophysical properties, revealing two major clusters matching FSI (green) and RSI (orange) subtypes. **B:** Decomposition of PC1 and PC2 loading by each intrinsic biophysical property.

**Figure S2.**
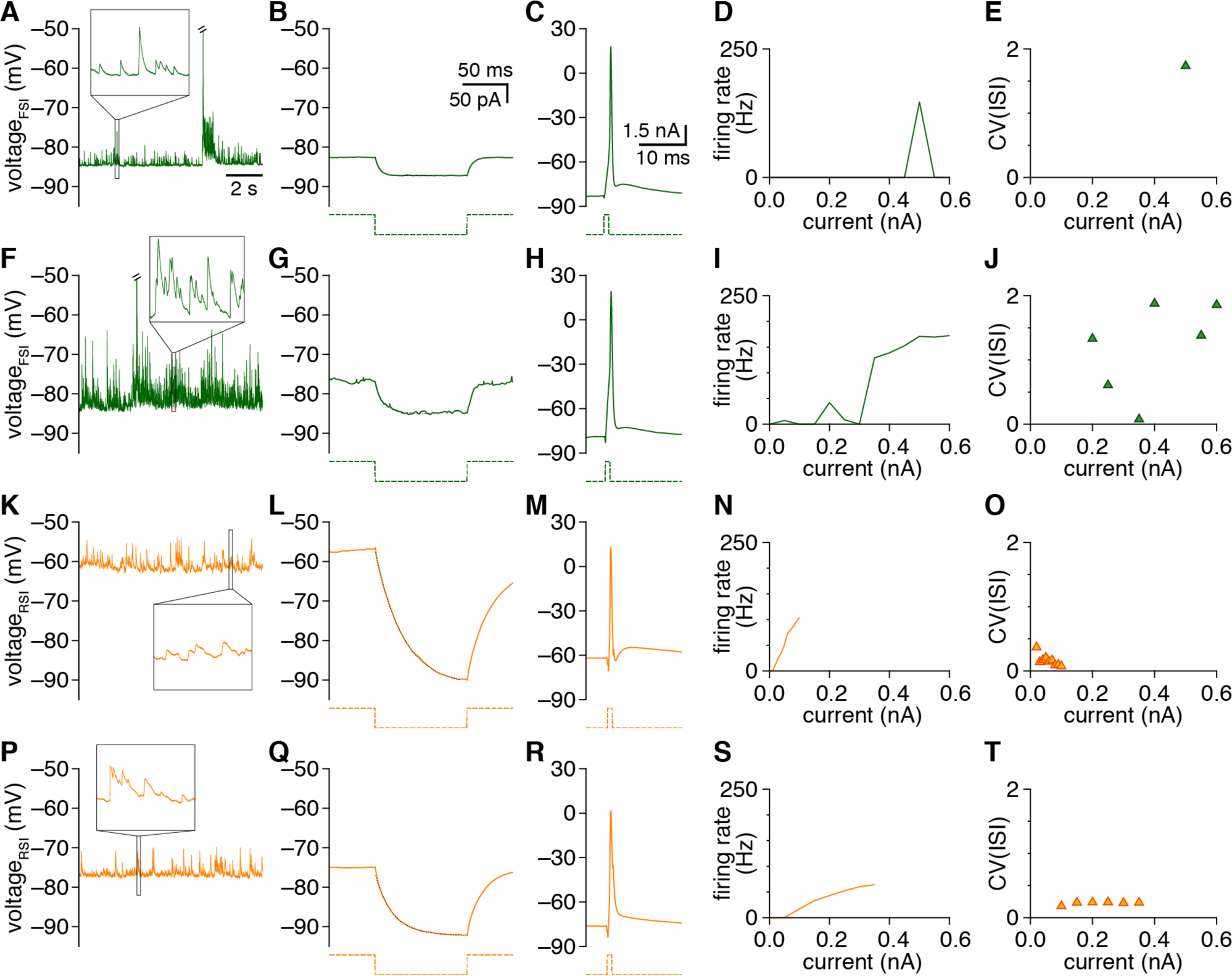
EPL-IN subtypes exhibit stark differences in most intrinsic biophysical properties. **A-E :** Diverse responses used to calculate intrinsic biophysical properties for the example FSI from Figure 1A, including: spontaneous activity at resting membrane potential (**A**), mean response to negative step current injection, with single-exponential fit (dashed black line) (**B**), mean spike waveform evoked by 1-ms suprathreshold current injection (**C**), firing rate-current relationship (**D**), and interspike interval (ISI) coefficient of variation evoked by positive step current injection (**E**). Spontaneous spike in **A** truncated to better visualize synaptic activity. Inset in **A**: magnification of spontaneous synaptic events. **F-T:** Same as **A-E** for the example FSI and RSIs from Figure 1C,E,G. Insets in **A,F,K,P** are identically scaled.

**Figure S3.**
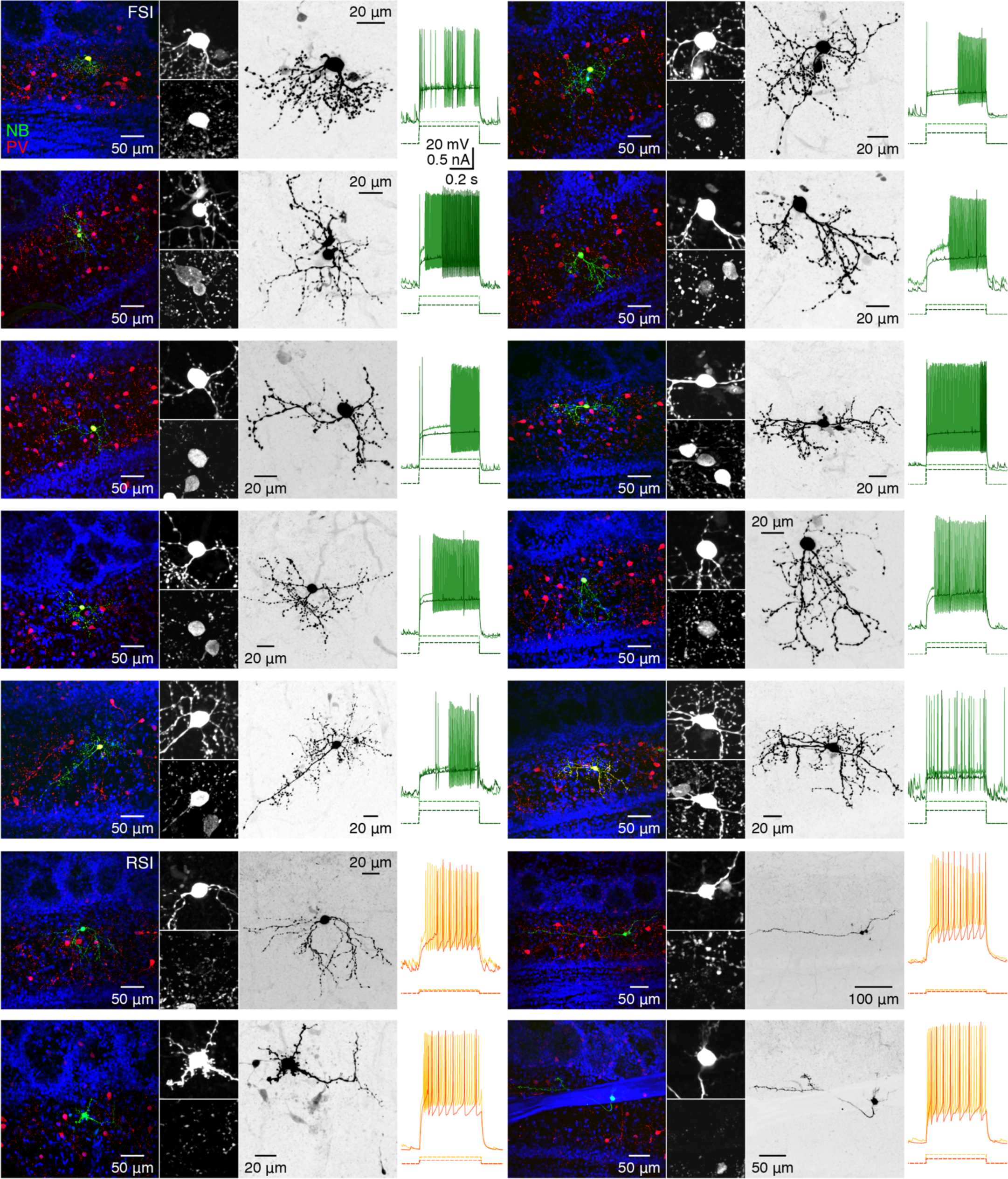
PV expression distinguishes FSIs from RSIs. Intracellular NB and post hoc PV staining with 50-μm magnified inset of somata (left), inverted NB (middle), and step current-evoked spiking (right) of a panel of EPL-INs. Spiking responses are color-coded to reflect FSI vs. RSI physiology, as in Figure 1.

**Figure S4.**
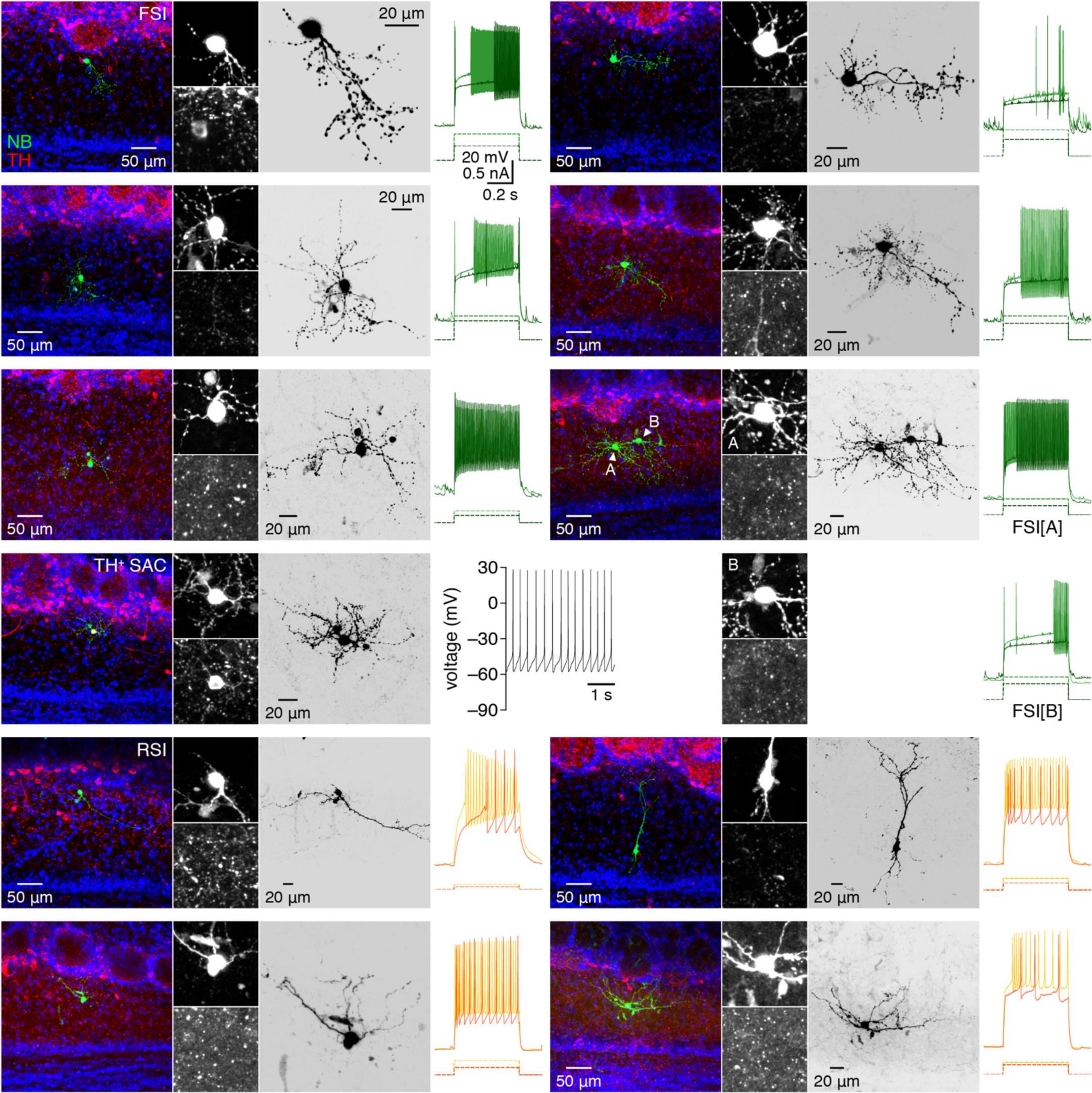
Neither FSIs nor RSIs are dopaminergic. Intracellular NB and post hoc TH staining with 50-μm magnified inset of somata (left), inverted NB (middle), and step current-evoked spiking (right) of a panel of EPL-INs. Spiking responses are color-coded to reflect FSI vs. RSI physiology, as in Figure 1. An example TH^+^ SAC exhibiting tonic spontaneous firing is additionally included as positive control for TH staining.

**Figure S5.**
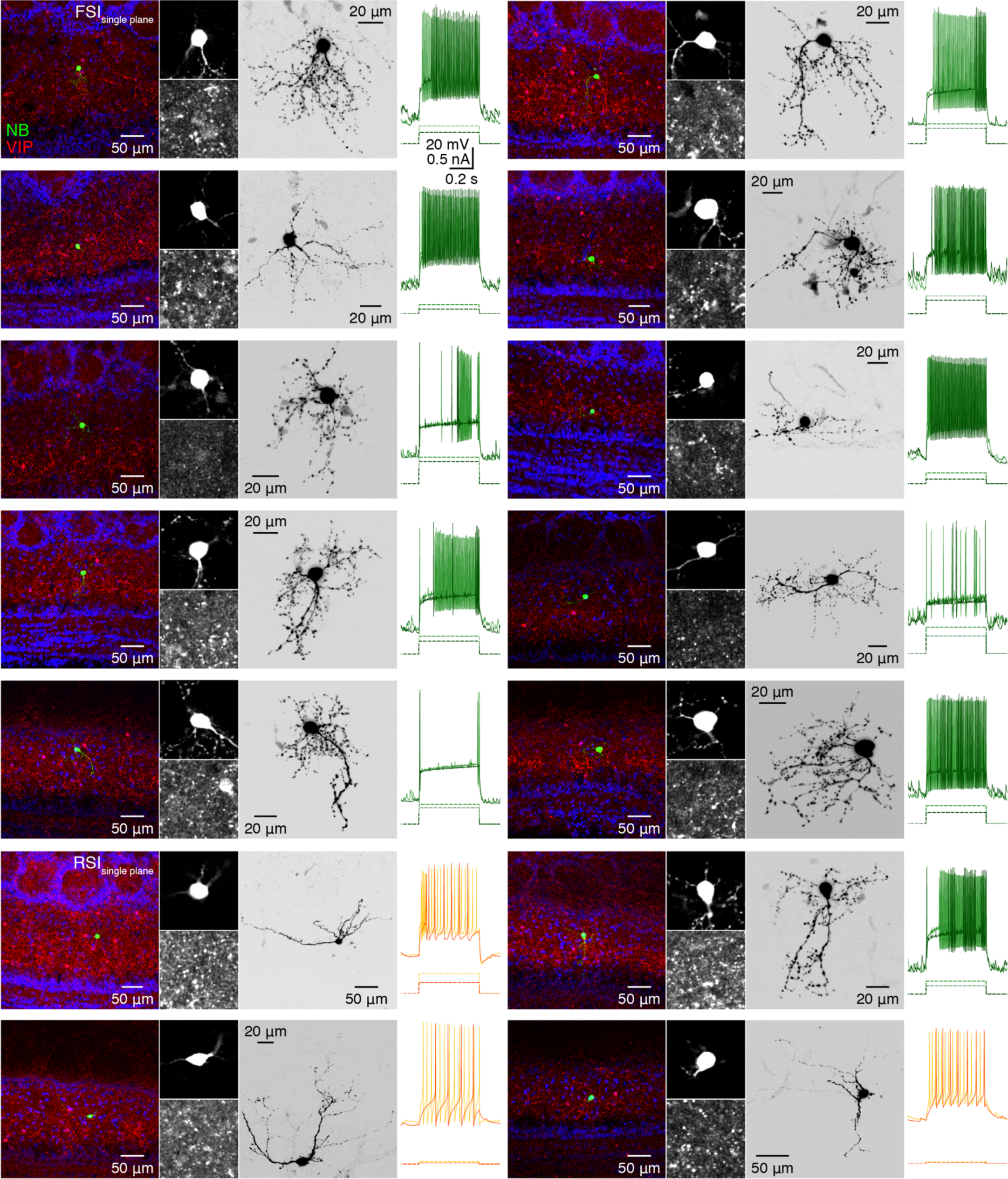
VIP expression poorly distinguishes FSIs and RSIs. Intracellular NB and post hoc VIP staining with 50-μm magnified inset of somata (left; single optical confocal planes), inverted NB (middle; maximum-intensity confocal projection), and step current-evoked spiking (right) of a panel of EPL-INs. Spiking responses are color-coded to reflect FSI vs. RSI physiology, as in Figure 1.

**Figure S6.**
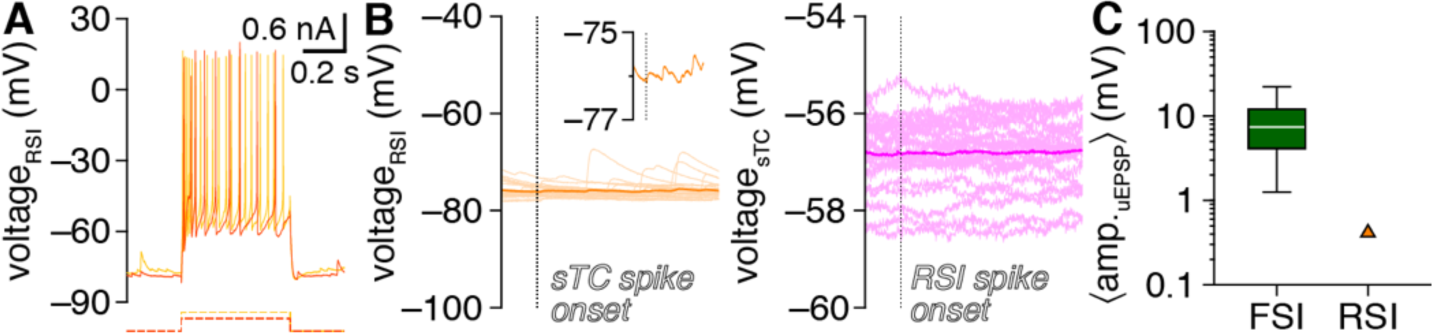
Unitary MTC-to-RSI excitation in a solitary example was distinctly weaker than FSI excitation. **A,B:** Step current-evoked spiking response (**A**) and unitary synaptic interactions (**B**) for the solitary MTC–RSI pair exhibiting significant unitary MTC-to-RSI excitation (morphology not recovered). Postsynaptic RSI voltage shown on same scale as Figure 1O,P for comparison to postsynaptic FSI responses. Inset: magnification of mean postsynaptic RSI voltage. **C:** The MTC-to-RSI uEPSP amplitude was markedly weaker than FSI uEPSPs (n=69).

**Figure S7.**
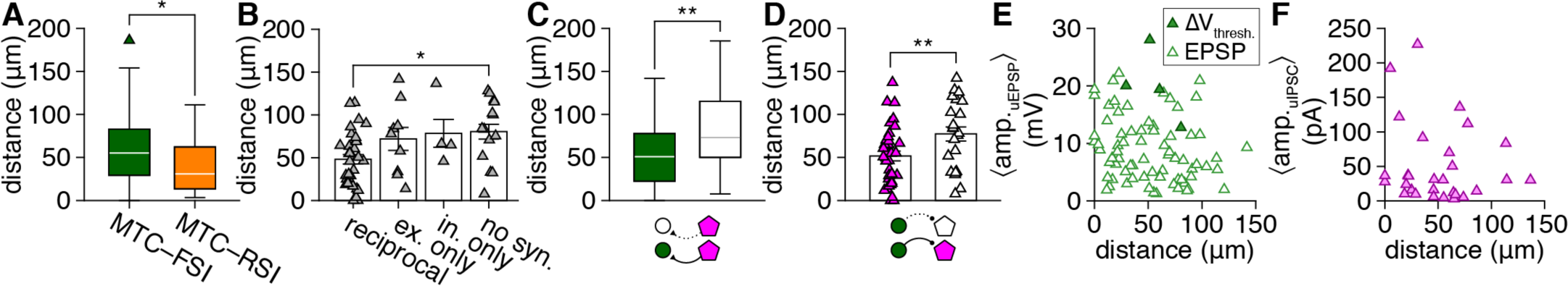
Connected MTC–FSI pairs exhibit shorter intersomatic distances than unconnected pairs. **A:** MTC–RSI pairs (n=19) exhibited modestly shorter intersomatic distances than MTC–FSI pairs (n=97) (*p=0.02, Wilcoxon rank-sum test). **B:** Among MTC–FSI pairs, reciprocally-connected pairs exhibited shorter intersomatic distances than unconnected pairs (p=0.02, F3,55=3.4, one-way ANOVA; reciprocal vs. excitation only: p=0.3, reciprocal vs. inhibition only: p=0.4, reciprocal vs. unconnected: p=0.03, excitation only vs. inhibition only: p=1.0, excitation only vs. unconnected: p=0.9, inhibition only vs. unconnected: p=1.0, post-hoc Tukey-Kramer test). **C:** MTC–FSI pairs with significant unitary MTC-to-FSI excitation exhibited shorter intersomatic distances than pairs with no excitatory connectivity (**p=2.5×10^−3^, Wilcoxon rank-sum test). **D:** MTC–FSI pairs with significant unitary FSI-to-MTC inhibition exhibited shorter intersomatic distances than pairs with no inhibitory connectivity (**p=9.7×10^−3^, t_57_=2.7, two-sample t-test). Analysis restricted to pairs with voltage-clamped MTCs (and therefore sensitive detection of unitary inhibition). **E,F:** Neither MTC-to-FSI uEPSP amplitudes (**E**) nor FSI-to-MTC uIPSC amplitudes (**F**) correlated with intersomatic distance (uEPSP: n=77 connections; p=0.1, t_75_=1.7, linear regression, slope not significantly different from 0; uIPSC: n=29 connections; p=0.4, t_27_=0.8, linear regression, slope not significantly different from 0). Pairs lacking connectivity (i.e., uEPSP or uIPSC amplitude of zero) not included in analysis.

**Figure S8.**
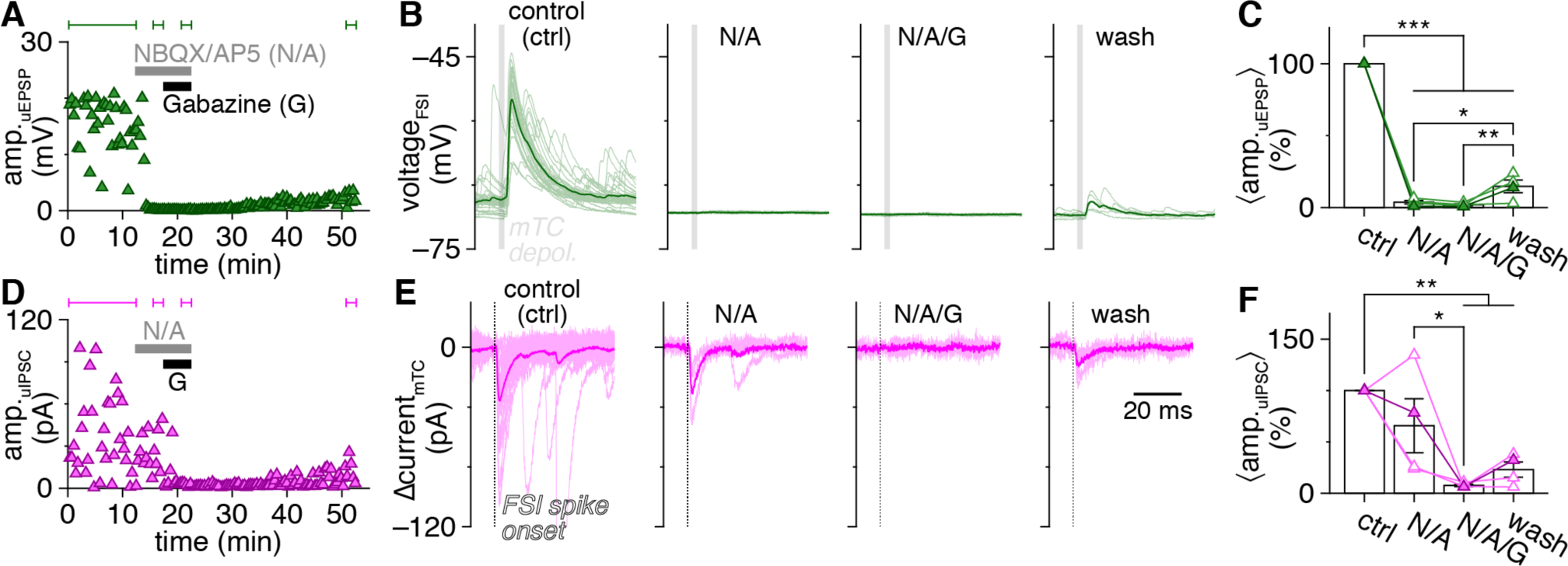
Unitary MTC–FSI synaptic pharmacology. **A:** Recording from an example MTC–FSI pair (morphology not recovered) showing MTC-to-FSI uEPSP amplitudes before and after combined bath application of glutamatergic antagonists NBQX (10 μM) and AP5 (50 μM) and subsequent application of GABA_A_R antagonist Gabazine (10 μM). **B:** Postsynaptic FSI voltages from the pair in **A**. Traces in each subplot correspond to the bracketed trials in **A**. **C:** Unitary MTC-to-FSI excitation was blocked by combined application of NBQX and AP5 and partially recovered upon wash-out in 4 MTC–FSI pairs (p=2.2×10^−12^, F_3,12_=414.9, one-way ANOVA; ctrl vs. N/A: ***p=5.5×10^−9^, ctrl vs. N/A/G: ***p=5.5×10^−9^, ctrl vs. wash: ***p=5.5×10^−9^, N/A vs. N/A/G: p=0.9, N/A vs. wash: *p=0.03, N/A/G vs. wash: **p=0.01, post-hoc Tukey-Kramer test). **D,E:** Same as **A,B** for FSI-to-MTC uIPSCs recorded in the same example pair. **F:** Unitary FSI-to-MTC inhibition was blocked by application of Gabazine and partially recovered upon wash-out in the same 4 MTC–FSI pairs as **C** (p=1.7×10^−3^, F_3,12_=9.5, one-way ANOVA; ctrl vs. N/A: p=0.33, ctrl vs. N/A/G: **p=2.1×10^−3^, ctrl vs. wash: **p=8.3×10^−3^, N/A vs. N/A/G: *p=0.046, N/A vs. wash: p=0.17, N/A/G vs. wash: p=0.85, post-hoc Tukey-Kramer test). Filled symbols in **C,F** correspond to the example pair shown.

**Figure S9.**
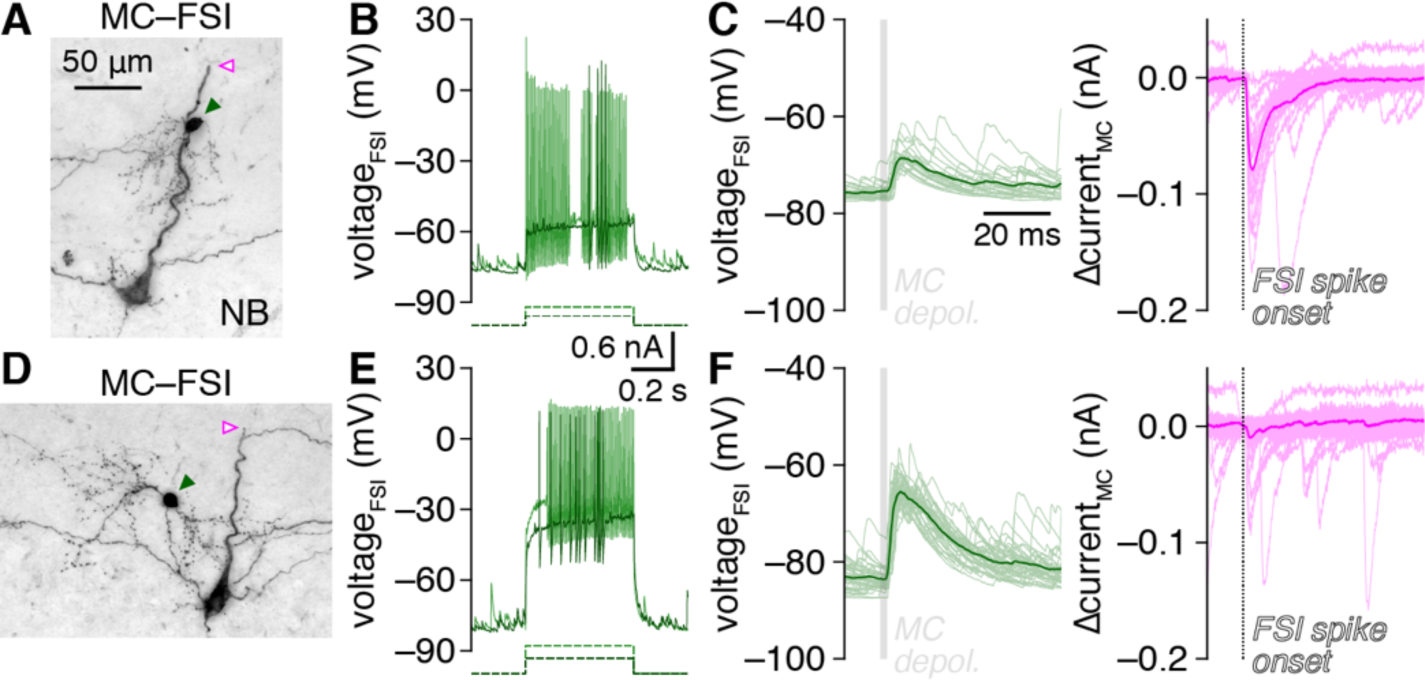
MTC–FSI connectivity is restricted to infraglomerular layers. **A:** Example MTC–FSI pair with MTC apical dendrite truncated prior to entering glomerular layer (open arrowhead). **B,C:** FSI fast-spiking response to step current injection (**B**) and unitary synaptic connectivity with MTC (**C**). **D-F:** Same as **A-C** for a second example MTC–FSI pair.

**Figure S10.**
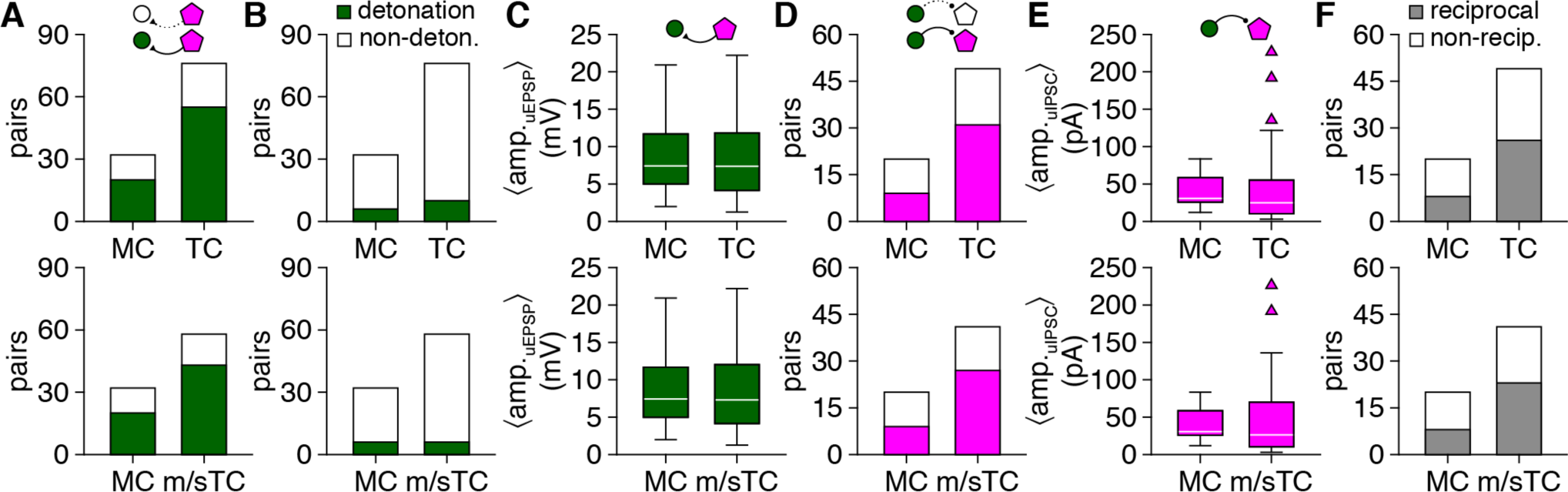
MCs and TCs exhibit similar unitary connectivity with FSIs. **A:** Detection of unitary FSI excitation did not significantly differ between MC–FSI and TC–FSI pairs (upper; p=0.3, χ^2^_[1]_=1.0, χ^2^ test) even when considering only middle and superficial TCs (m/sTCs) to exclude potential misclassification of deep TCs (lower; p=0.2, χ^2^_[1]_=1.3, χ^2^ test). **B:** The proportion of FSIs responding to unitary MTC release with detonation did not significantly differ between MC–FSI and TC–FSI pairs (upper; p=0.5, χ^2^_[1]_=0.6, χ^2^ test) or between MC–FSI and m/sTC–FSI pairs (lower; p=0.3, χ^2^_[1]_=1.3, χ^2^ test). **C:** FSI uEPSP amplitudes did not significantly differ between MC-FSI and TC-FSI pairs (upper; p=1.0, Wilcoxon rank-sum test) or between MC–FSI and m/sTC–FSI pairs (lower, p=1.0, Wilcoxon rank-sum test). **D:** Detection of unitary MTC inhibition did not significantly differ between MC–FSI and TC–FSI pairs (upper; p=0.2, χ^2^_[1]_=1.9, χ^2^ test) or between MC–FSI and m/sTC–FSI pairs (lower; p=0.1, χ^2^_[1]_=2.4, χ^2^ test); only voltage-clamped MTCs were considered for peak detection sensitivity. **E:** MTC uIPSC amplitudes did not significantly differ between MC–FSI and TC–FSI pairs (upper; p=0.6, Wilcoxon rank-sum test) or between MC–FSI and m/sTC–FSI pairs (lower; p=0.6, Wilcoxon rank-sum test). **F:** The proportion of MTC–FSI pairs exhibiting reciprocal unitary connectivity did not differ between MC–FSI and TC–FSI pairs (upper; p=0.3, χ^2^_[1]_=1.0, χ^2^ test) or between MC–FSI and m/sTC–FSI pairs (lower; p=0.2, χ^2^_[1]_=1.4, χ^2^ test).

**Figure S11.**
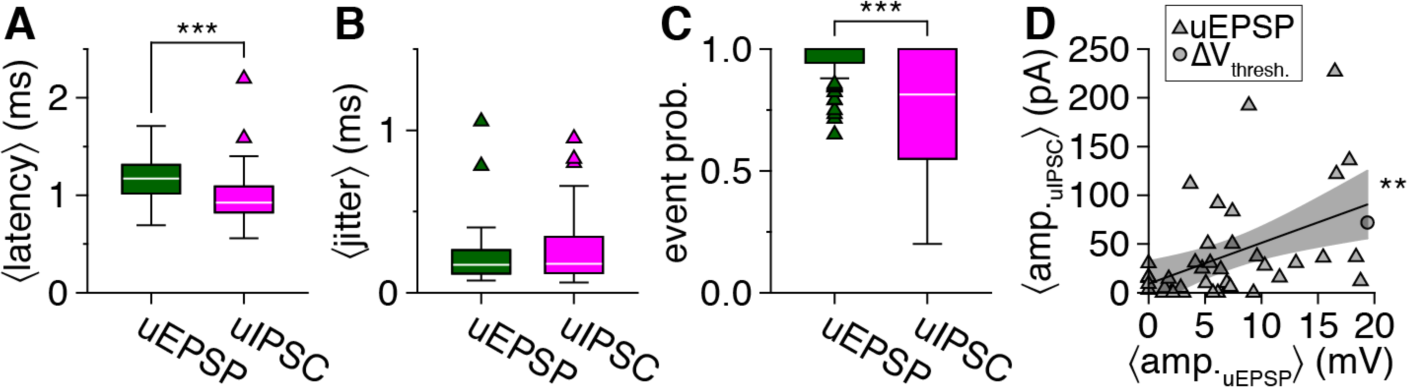
Comparison of unitary FSI-to-MTC and MTC-to-FSI synaptic transmission properties. **A:** FSI-to-MTC uIPSC latency (n=44 connections) was significantly shorter than MTC-to-FSI uEPSP latency (n=31 connections) (***p=8.0×10^−5^, Wilcoxon rank-sum test). **B:** FSI-to-MTC uIPSC jitter (n=44 connections) and MTC-to-FSI uEPSP jitter (n=31 connections) were equivalent (p=0.5, Wilcoxon rank-sum test). **C:** Trial-to-trial FSI-to-MTC uIPSC event probability (n=44 connections) was significantly lower than trial-to-trial MTC-to-FSI uEPSP event probability (n=79 connections) (***p=5.4×10^−8^, Wilcoxon rank-sum test). Unitary FSI-to-MTC IPSP latency, jitter, and probability not included in comparisons due to limited unitary IPSP detection sensitivity (Figure 1U). **D:** Across all MTC–FSI pairs with at least one direction of unitary connectivity, FSI-to-MTC uIPSC amplitude positively correlated with MTC-to-FSI uEPSP amplitude (n=41 pairs; **p=1.9×10^−3^, t_39_=3.3, R^2^=0.22, linear regression, slope significantly different from 0). For pairs exhibiting exclusive FSI detonation, uEPSP amplitudes were estimated as the difference between resting membrane potential and spike threshold (ΔV_thresh._), as in Figure 3G. Shading denotes 95% confidence interval.

**Figure S12.**
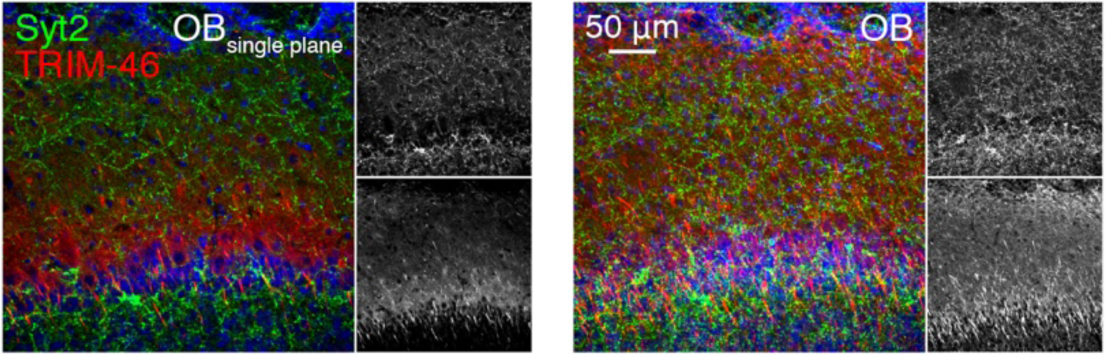
Syt2 clusters do not selectively target MTC axon initial segments. Single confocal optical plane (left) and maximum-intensity projection (right; ∼50 μm depth) of Syt2 and axon initial segment component TRIM-46 in the OB, revealing an absence of clear Chandelier-like innervation of MTCs.

**Figure S13.**
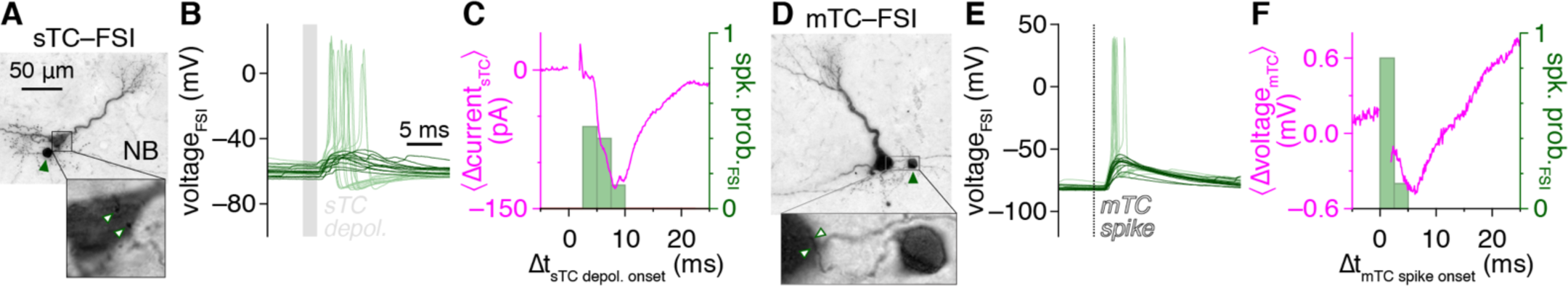
Comparison of spontaneously alternating FSI detonation vs. uEPSP trials reveals high-fidelity recurrent MTC inhibition. **A,B:** Example MTC–FSI pair (**A**) in which unitary MTC release triggers FSI detonation on some trials (light green) and uEPSPs on other trials (dark green) (**B**). **C:** Subtraction of mean MTC currents across FSI detonation vs. uEPSP trials from **B** isolates IPSC waveforms time-locked to FSI detonation. **D-F:** Same as **A-C** for an example MTC-FSI pair recorded in current-clamp, revealing isolation of an IPSP waveform time-locked to FSI detonation.

## Funding

This work was supported by National Institute on Deafness and Other Communication Disorders (https://www.nidcd.nih.gov/) grants R01DC016560 to N.N.U. and R01DC021296 to S.D.B. The funders had no role in study design, data collection and analysis, decision to publish, or preparation of the manuscript.

## Acknowledgments

We thank members of the Cheetham and Haas laboratories for helpful discussion.

## Author contributions

S.D.B. and N.N.U. designed experiments; S.D.B. and C.M.M. performed experiments; S.D.B., C.M.M., and N.N.U. analyzed data and wrote the paper.

## Competing interests

The authors declare no competing interests.

## Notes

### Competing Interest Statement

The authors have declared no competing interest.

## References

Aghvami SS, Kubota Y, Egger V (2022) Anatomical and Functional Connectivity at the Dendrodendritic Reciprocal Mitral Cell–Granule Cell Synapse: Impact on Recurrent and Lateral Inhibition. Front Neuroanat 16:1–18.

Arneodo EM, Penikis KB, Rabinowitz N, Licata A, Cichy A, Zhang J, Bozza T, Rinberg D (2018) Stimulus dependent diversity and stereotypy in the output of an olfactory functional unit. Nat Commun 9:1347.

Arshadi C, Günther U, Eddison M, Harrington KIS, Ferreira TA (2021) SNT: a unifying toolbox for quantification of neuronal anatomy. Nat Meth 18:374–377.

Batista-Brito R, Close J, Machold R, Fishell G (2008) The distinct temporal origins of olfactory bulb interneuron subtypes. J Neurosci 28:3966–3975.

Borst JG, Helmchen F, Sakmann B (1995) Pre- and postsynaptic whole-cell recordings in the medial nucleus of the trapezoid body of the rat. J Physiol (Lond) 489 (Pt 3):825–840.

Brunjes PC, Feldman S, Osterberg SK (2016) The Pig Olfactory Brain: A Primer. Chem Senses 41:415–425.

Buhl EH, Cobb SR, Halasy K, Somogyi P (1995) Properties of unitary IPSPs evoked by anatomically identified basket cells in the rat hippocampus. Eur J Neurosci 7:1989–2004.

Burton SD (2017) Inhibitory circuits of the mammalian main olfactory bulb. J Neurophysiol 118:2034–2051.

Burton SD, Brown A, Eiting TP, Youngstrom IA, Rust TC, Schmuker M, Wachowiak M (2022) Mapping odorant sensitivities reveals a sparse but structured representation of olfactory chemical space by sensory input to the mouse olfactory bulb. eLife 11.

Burton SD, Ermentrout GB, Urban NN (2012) Intrinsic heterogeneity in oscillatory dynamics limits correlation-induced neural synchronization. J Neurophysiol 108:2115–2133.

Burton SD, Larocca G, Liu A, Cheetham CEJ, Urban NN (2017) Olfactory Bulb Deep Short-Axon Cells Mediate Widespread Inhibition of Tufted Cell Apical Dendrites. J Neurosci 37:1117– 1138.

Burton SD, Urban NN (2014) Greater excitability and firing irregularity of tufted cells underlies distinct afferent-evoked activity of olfactory bulb mitral and tufted cells. J Physiol (Lond) 592:2097–2118.

Burton SD, Urban NN (2015) Rapid Feedforward Inhibition and Asynchronous Excitation Regulate Granule Cell Activity in the Mammalian Main Olfactory Bulb. J Neurosci 35:14103–14122.

Burton SD, Urban NN (2021) Cell and circuit origins of fast network oscillations in the mammalian main olfactory bulb. eLife 10.

Campanac E, Gasselin C, Baude A, Rama S, Ankri N, Debanne D (2013) Enhanced intrinsic excitability in basket cells maintains excitatory-inhibitory balance in hippocampal circuits. Neuron 77:712–722.

Carlson GC, Shipley MT, Keller A (2000) Long-lasting depolarizations in mitral cells of the rat olfactory bulb. J Neurosci 20:2011–2021.

Chae H, Banerjee A, Dussauze M, Albeanu DF (2022) Long-range functional loops in the mouse olfactory system and their roles in computing odor identity. Neuron.

Compans B, Burrone J (2023) Chandelier cells shine a light on the formation of GABAergic synapses. Curr Opin Neurobiol 80:102697.

Crespo C, Blasco-Ibáñez JM, Marqués-Marí AI, Alonso JR, Briñón JG, Martínez-Guijarro FJ (2002) Vasoactive intestinal polypeptide-containing elements in the olfactory bulb of the hedgehog (Erinaceus europaeus). J Chem Neuroanat 24:49–63.

Crespo C, Liberia T, Blasco-Ibáñez JM, Nácher J, Varea E (2013) The circuits of the olfactory bulb. The exception as a rule. Anat Rec (Hoboken) 296:1401–1412.

Darcy DP, Isaacson JS (2010) Calcium-permeable AMPA receptors mediate glutamatergic signaling in neural precursor cells of the postnatal olfactory bulb. J Neurophysiol 103:1431– 1437.

De Saint Jan D, Hirnet D, Westbrook GL, Charpak S (2009) External tufted cells drive the output of olfactory bulb glomeruli. J Neurosci 29:2043–2052.

Dietz SB, Markopoulos F, Murthy VN (2011) Postnatal development of dendrodendritic inhibition in the Mammalian olfactory bulb. Front Cell Neurosci 5:10.

Eccles JC, Llinaś R, Sasaki K (1966) The excitatory synaptic action of climbing fibres on the Purkinje cells of the cerebellum. J Physiol (Lond) 182:268–296.

Egger V, Kuner T (2021) Olfactory bulb granule cells: specialized to link coactive glomerular columns for percept generation and discrimination of odors. Cell Tissue Res 383:495–506.

Egger V, Urban NN (2006) Dynamic connectivity in the mitral cell-granule cell microcircuit. Semin Cell Dev Biol 17:424–432.

Eyre MD, Renzi M, Farrant M, Nusser Z (2012) Setting the time course of inhibitory synaptic currents by mixing multiple GABA(A) receptor α subunit isoforms. J Neurosci 32:5853–5867.

Freund TF, Katona I (2007) Perisomatic inhibition. Neuron 56:33–42.

Fritschy JM, Mohler H (1995) GABAA-receptor heterogeneity in the adult rat brain: differential regional and cellular distribution of seven major subunits. J Comp Neurol 359:154–194.

Fukunaga I, Berning M, Kollo M, Schmaltz A, Schaefer AT (2012) Two distinct channels of olfactory bulb output. Neuron 75:320–329.

Galán RF, Fourcaud-Trocmé N, Ermentrout GB, Urban NN (2006) Correlation-induced synchronization of oscillations in olfactory bulb neurons. J Neurosci 26:3646–3655.

Garcia I, Bhullar PK, Tepe B, Ortiz-Guzman J, Huang L, Herman AM, Chaboub L, Deneen B, Justice NJ, Arenkiel BR (2016) Local corticotropin releasing hormone (CRH) signals to its receptor CRHR1 during postnatal development of the mouse olfactory bulb. Brain Struct Funct 221:1–20.

Geramita MA, Burton SD, Urban NN (2016) Distinct lateral inhibitory circuits drive parallel processing of sensory information in the mammalian olfactory bulb. eLife 5.

Giustetto M, Bovolin P, Fasolo A, Bonino M, Cantino D, Sassoe-Pognetto M (1997) Glutamate receptors in the olfactory bulb synaptic circuitry: heterogeneity and synaptic localization of N-methyl-D-aspartate receptor subunit 1 and AMPA receptor subunit 1. Neuroscience 76:787– 798.

Gödde K, Gschwend O, Puchkov D, Pfeffer CK, Carleton A, Jentsch TJ (2016) Disruption of Kcc2-dependent inhibition of olfactory bulb output neurons suggests its importance in odour discrimination. Nat Commun 7:12043.

Haddad R, Lanjuin A, Madisen L, Zeng H, Murthy VN, Uchida N (2013) Olfactory cortical neurons read out a relative time code in the olfactory bulb. Nat Neurosci 16:949–957.

Hamilton KA, Parrish-Aungst S, Margolis FL, Erdelyi F, Szabo G, Puche AC (2008) Sensory deafferentation transsynaptically alters neuronal GluR1 expression in the external plexiform layer of the adult mouse main olfactory bulb. Chem Senses 33:201–210.

Hefft S, Jonas P (2005) Asynchronous GABA release generates long-lasting inhibition at a hippocampal interneuron-principal neuron synapse. Nat Neurosci 8:1319–1328.

Hippenmeyer S, Vrieseling E, Sigrist M, Portmann T, Laengle C, Ladle DR, Arber S (2005) A developmental switch in the response of DRG neurons to ETS transcription factor signaling. PLoS Biol 3:e159.

Hooks BM, Lin JY, Guo C, Svoboda K (2015) Dual-channel circuit mapping reveals sensorimotor convergence in the primary motor cortex. J Neurosci 35:4418–4426.

Hu H, Gan J, Jonas P (2014) Interneurons. Fast-spiking, parvalbumin⁺ GABAergic interneurons: from cellular design to microcircuit function. Science 345:1255263.

Huang L, Garcia I, Jen H-I, Arenkiel BR (2013) Reciprocal connectivity between mitral cells and external plexiform layer interneurons in the mouse olfactory bulb. Front Neural Circuits 7:32.

Igarashi KM, Ieki N, An M, Yamaguchi Y, Nagayama S, Kobayakawa K, Kobayakawa R, Tanifuji M, Sakano H, Chen WR, Mori K (2012) Parallel mitral and tufted cell pathways route distinct odor information to different targets in the olfactory cortex. J Neurosci 32:7970–7985.

Isaacson JS (2001) Mechanisms governing dendritic gamma-aminobutyric acid (GABA) release in the rat olfactory bulb. Proc Natl Acad Sci USA 98:337–342.

Isaacson JS, Strowbridge BW (1998) Olfactory reciprocal synapses: dendritic signaling in the CNS. Neuron 20:749–761.

Joye DAM, Rohr KE, Keller D, Inda T, Telega A, Pancholi H, Carmona-Alcocer V, Evans JA (2020) Reduced VIP Expression Affects Circadian Clock Function in VIP-IRES-CRE Mice (JAX 010908). J Biol Rhythms 35:340–352.

Kapoor V, Urban NN (2006) Glomerulus-Specific, Long-Latency Activity in the Olfactory Bulb Granule Cell Network. J Neurosci 26:11709–11719.

Kashiwadani H, Sasaki YF, Uchida N, Mori K (1999) Synchronized oscillatory discharges of mitral/tufted cells with different molecular receptive ranges in the rabbit olfactory bulb. J Neurophysiol 82:1786–1792.

Kato HK, Gillet SN, Peters AJ, Isaacson JS, Komiyama T (2013) Parvalbumin-expressing interneurons linearly control olfactory bulb output. Neuron 80:1218–1231.

Kay LM (2014) Circuit oscillations in odor perception and memory. Prog Brain Res 208:223–251.

Kosaka K, Heizmann CW, Kosaka T (1994) Calcium-binding protein parvalbumin-immunoreactive neurons in the rat olfactory bulb. 2. Postnatal development. Exp Brain Res 99:205–213.

Kosaka T, Komada M, Kosaka K (2008) Sodium channel cluster, betaIV-spectrin and ankyrinG positive “hot spots” on dendritic segments of parvalbumin-containing neurons and some other neurons in the mouse and rat main olfactory bulbs. Neurosci Res 62:176–186.

Kosaka T, Kosaka K (2008) Heterogeneity of parvalbumin-containing neurons in the mouse main olfactory bulb, with special reference to short-axon cells and betaIV-spectrin positive dendritic segments. Neurosci Res 60:56–72.

Kosaka T, Kosaka K (2010) Heterogeneity of calbindin-containing neurons in the mouse main olfactory bulb: I. General description. Neurosci Res 67:275–292.

Kosaka T, Pignatelli A, Kosaka K (2020) Heterogeneity of tyrosine hydroxylase expressing neurons in the main olfactory bulb of the mouse. Neurosci Res 157:15–33.

Lage-Rupprecht V, Zhou L, Bianchini G, Aghvami SS, Mueller M, Rózsa B, Sassoè-Pognetto M, Egger V (2020) Presynaptic NMDARs cooperate with local spikes toward GABA release from the reciprocal olfactory bulb granule cell spine. eLife 9.

Lagier S, Panzanelli P, Russo RE, Nissant A, Bathellier B, Sassoè-Pognetto M, Fritschy J-M, Lledo P-M (2007) GABAergic inhibition at dendrodendritic synapses tunes gamma oscillations in the olfactory bulb. Proc Natl Acad Sci USA 104:7259–7264.

Lepousez G, Csaba Z, Bernard V, Loudes C, Videau C, Lacombe J, Epelbaum J, Viollet C (2010) Somatostatin interneurons delineate the inner part of the external plexiform layer in the mouse main olfactory bulb. J Comp Neurol 518:1976–1994.

Lepousez G, Lledo P-M (2013) Odor discrimination requires proper olfactory fast oscillations in awake mice. Neuron 80:1010–1024.

Li G, Cleland TA (2017) A coupled-oscillator model of olfactory bulb gamma oscillations. PLoS Comput Biol 13:e1005760.

Li J, Ishii T, Feinstein P, Mombaerts P (2004) Odorant receptor gene choice is reset by nuclear transfer from mouse olfactory sensory neurons. Nature 428:393–399.

Liberia T, Blasco-Ibáñez JM, Nácher J, Varea E, Zwafink V, Crespo C (2012) Characterization of a population of tyrosine hydroxylase-containing interneurons in the external plexiform layer of the rat olfactory bulb. Neuroscience 217:140–153.

Lindeman S, Fu X, Reinert JK, Fukunaga I (2024) Value-related learning in the olfactory bulb occurs through pathway-dependent perisomatic inhibition of mitral cells. PLoS Biol 22:e3002536.

Liu G, Froudarakis E, Patel JM, Kochukov MY, Pekarek B, Hunt PJ, Patel M, Ung K, Fu C-H, Jo J, Lee H-K, Tolias AS, Arenkiel BR (2019) Target specific functions of EPL interneurons in olfactory circuits. Nat Commun 10:1–14.

López-Mascaraque L, De Carlos JA, Valverde F (1990) Structure of the olfactory bulb of the hedgehog (Erinaceus europaeus): a Golgi study of the intrinsic organization of the superficial layers. J Comp Neurol 301:243–261.

Ma J, Lowe G (2007) Calcium permeable AMPA receptors and autoreceptors in external tufted cells of rat olfactory bulb. Neuroscience 144:1094–1108.

Maccaferri G, Lacaille J-C (2003) Interneuron Diversity series: Hippocampal interneuron classifications--making things as simple as possible, not simpler. Trends Neurosci 26:564– 571.

Madisen L, Zwingman TA, Sunkin SM, Oh SW, Zariwala HA, Gu H, Ng LL, Palmiter RD, Hawrylycz MJ, Jones AR, Lein ES, Zeng H (2010) A robust and high-throughput Cre reporting and characterization system for the whole mouse brain. Nat Neurosci 13:133–140.

Maher BJ, McGinley MJ, Westbrook GL (2009) Experience-dependent maturation of the glomerular microcircuit. Proc Natl Acad Sci USA 106:16865–16870.

Marín O (2012) Interneuron dysfunction in psychiatric disorders. 13:107–120.

Markram H, Toledo-Rodriguez M, Wang Y, Gupta A, Silberberg G, Wu C (2004) Interneurons of the neocortical inhibitory system. 5:793–807.

McIntyre ABR, Cleland TA (2016) Biophysical constraints on lateral inhibition in the olfactory bulb. J Neurophysiol 115:2937–2949.

McNaughton BL, Morris RGM (1987) Hippocampal synaptic enhancement and information storage within a distributed memory system. Trends Neurosci 10:408–415.

McTavish TS, Migliore M, Shepherd GM, Hines ML (2012) Mitral cell spike synchrony modulated by dendrodendritic synapse location. Front Comput Neurosci 6:3.

Molnár G, Oláh S, Komlósi G, Füle M, Szabadics J, Varga C, Barzó P, Tamás G (2008) Complex events initiated by individual spikes in the human cerebral cortex. PLoS Biol 6:e222.

Montague AA, Greer CA (1999) Differential distribution of ionotropic glutamate receptor subunits in the rat olfactory bulb. J Comp Neurol 405:233–246.

Mueller M, Egger V (2020) Dendritic integration in olfactory bulb granule cells upon simultaneous multispine activation: Low thresholds for nonlocal spiking activity. PLoS Biol 18:e3000873.

Nagayama S, Homma R, Imamura F (2014) Neuronal organization of olfactory bulb circuits. Front Neural Circuits 8:98.

Najac M, Sanz Diez A, Kumar A, Benito N, Charpak S, De Saint Jan D (2015) Intraglomerular lateral inhibition promotes spike timing variability in principal neurons of the olfactory bulb. J Neurosci 35:4319–4331.

Nigro MJ, Kirikae H, Kjelsberg K, Nair RR, Witter MP (2021) Not All That Is Gold Glitters: PV-IRES-Cre Mouse Line Shows Low Efficiency of Labeling of Parvalbumin Interneurons in the Perirhinal Cortex. Front Neural Circuits 15:781928.

Ona-Jodar T, Lage-Rupprecht V, Abraham NM, Rose CR, Egger V (2020) Local Postsynaptic Signaling on Slow Time Scales in Reciprocal Olfactory Bulb Granule Cell Spines Matches Asynchronous Release. Front Synaptic Neurosci 12:551691.

Pang ZP, Melicoff E, Padgett D, Liu Y, Teich AF, Dickey BF, Lin W, Adachi R, Südhof TC (2006) Synaptotagmin-2 is essential for survival and contributes to Ca2+ triggering of neurotransmitter release in central and neuromuscular synapses. J Neurosci 26:13493– 13504.

Panzanelli P, Perazzini A-Z, Fritschy J-M, Sassoè-Pognetto M (2005) Heterogeneity of gamma-aminobutyric acid type A receptors in mitral and tufted cells of the rat main olfactory bulb. J Comp Neurol 484:121–131.

Petralia RS, Wang YX, Mayat E, Wenthold RJ (1997) Glutamate receptor subunit 2-selective antibody shows a differential distribution of calcium-impermeable AMPA receptors among populations of neurons. J Comp Neurol 385:456–476.

Petralia RS, Wenthold RJ (1992) Light and electron immunocytochemical localization of AMPA-selective glutamate receptors in the rat brain. J Comp Neurol 318:329–354.

Pignatelli A, Ackman JB, Vigetti D, Beltrami AP, Zucchini S, Belluzzi O (2009) A potential reservoir of immature dopaminergic replacement neurons in the adult mammalian olfactory bulb. Pflugers Arch 457:899–915.

Pirker S, Schwarzer C, Wieselthaler A, Sieghart W, Sperk G (2000) GABA(A) receptors: immunocytochemical distribution of 13 subunits in the adult rat brain. Neuroscience 101:815– 850.

Pressler RT, Strowbridge BW (2017) Direct Recording of Dendrodendritic Excitation in the Olfactory Bulb: Divergent Properties of Local and External Glutamatergic Inputs Govern Synaptic Integration in Granule Cells. J Neurosci 37:11774–11788.

Pressler RT, Strowbridge BW (2020) Activation of Granule Cell Interneurons by Two Divergent Local Circuit Pathways in the Rat Olfactory Bulb. J Neurosci 40:9701–9714.

Rall W (1967) Distinguishing theoretical synaptic potentials computed for different soma-dendritic distributions of synaptic input. J Neurophysiol 30:1138–1168.

Rall W, Shepherd GM (1968) Theoretical reconstruction of field potentials and dendrodendritic synaptic interactions in olfactory bulb. J Neurophysiol 31:884–915.

Rall W, Shepherd GM, Reese TS, Brightman MW (1966) Dendrodendritic synaptic pathway for inhibition in the olfactory bulb. Exp Neurol 14:44–56.

Schneider SP, Macrides F (1978) Laminar distributions of internuerons in the main olfactory bulb of the adult hamster. Brain Res Bull 3:73–82.

Schoppa NE (2006) Synchronization of olfactory bulb mitral cells by precisely timed inhibitory inputs. Neuron 49:271–283.

Schoppa NE, Urban NN (2003) Dendritic processing within olfactory bulb circuits. Trends Neurosci 26:501–506.

Shepherd GM, Chen WR, Greer CA (2004) Olfactory Bulb. In: The Synaptic Organization of the Brain, 5 ed. (Shepherd GM, ed), pp 159–204.

Silver RA (2010) Neuronal arithmetic. 11:474–489.

Smear M, Resulaj A, Zhang J, Bozza T, Rinberg D (2013) Multiple perceptible signals from a single olfactory glomerulus. Nat Neurosci 16:1687–1691.

Smith TC, Jahr CE (2002) Self-inhibition of olfactory bulb neurons. Nat Neurosci 5:760–766.

Sommeijer J-P, Levelt CN (2012) Synaptotagmin-2 is a reliable marker for parvalbumin positive inhibitory boutons in the mouse visual cortex. PLoS ONE 7:e35323.

Strowbridge BW (2009) Role of cortical feedback in regulating inhibitory microcircuits. Ann N Y Acad Sci 1170:270–274.

Tan J, Savigner A, Ma M, Luo M (2010) Odor information processing by the olfactory bulb analyzed in gene-targeted mice. Neuron 65:912–926.

Thomson AM, West DC, Hahn J, Deuchars J (1996) Single axon IPSPs elicited in pyramidal cells by three classes of interneurones in slices of rat neocortex. J Physiol (Lond) 496 (Pt 1):81–102.

Tiesinga P, Fellous J-M, Sejnowski TJ (2008) Regulation of spike timing in visual cortical circuits. 9:97–107.

Toida K, Kosaka K, Heizmann CW, Kosaka T (1994) Synaptic contacts between mitral/tufted cells and GABAergic neurons containing calcium-binding protein parvalbumin in the rat olfactory bulb, with special reference to reciprocal synapses between them. Brain Res 650:347–352.

Toida K, Kosaka K, Heizmann CW, Kosaka T (1996) Electron microscopic serial-sectioning/reconstruction study of parvalbumin-containing neurons in the external plexiform layer of the rat olfactory bulb. Neuroscience 72:449–466.

Trimble WS, Gray TS, Elferink LA, Wilson MC, Scheller RH (1990) Distinct patterns of expression of two VAMP genes within the rat brain. J Neurosci 10:1380–1387.

Urban NN, Castro JB (2010) Functional polarity in neurons: what can we learn from studying an exception? Curr Opin Neurobiol 20:538–542.

Urban NN, Sakmann B (2002) Reciprocal intraglomerular excitation and intra- and interglomerular lateral inhibition between mouse olfactory bulb mitral cells. J Physiol (Lond) 542:355–367.

Van Gehuchten A, Martin I (1891) Le bulbe olfactif chez quelques mammiferes. In: La Cellule, pp 205–237.

Vandael D, Jonas P (2024) Structure, biophysics, and circuit function of a “giant” cortical presynaptic terminal. Science 383:eadg6757.

Viollet C, Simon A, Tolle V, Labarthe A, Grouselle D, Loe-Mie Y, Simonneau M, Martel G, Epelbaum J (2017) Somatostatin-IRES-Cre Mice: Between Knockout and Wild-Type? Front Endocrinol (Lausanne) 8:131.

Vuong CK, Wei W, Lee J-A, Lin C-H, Damianov A, la Torre-Ubieta de L, Halabi R, Otis KO, Martin KC, O’Dell TJ, Black DL (2018) Rbfox1 Regulates Synaptic Transmission through the Inhibitory Neuron-Specific vSNARE Vamp1. Neuron 98:127–141.e127.

Wachowiak M (2011) All in a sniff: olfaction as a model for active sensing. Neuron 71:962–973.

Wang D, Wu J, Liu P, Li X, Li J, He M, Li A (2022) VIP interneurons regulate olfactory bulb output and contribute to odor detection and discrimination. Cell Rep 38:110383.

Wang Y, Hu P, Shan Q, Huang C, Huang Z, Chen P, Li A, Gong H, Zhou J-N (2021) Single-cell morphological characterization of CRH neurons throughout the whole mouse brain. BMC Biol 19:47.

Yokoi M, Mori K, Nakanishi S (1995) Refinement of odor molecule tuning by dendrodendritic synaptic inhibition in the olfactory bulb. Proc Natl Acad Sci USA 92:3371–3375.

Yu Y, Burton SD, Tripathy SJ, Urban NN (2015) Postnatal development attunes olfactory bulb mitral cells to high-frequency signaling. J Neurophysiol 114:2830–2842.

Zak JD, Whitesell JD, Schoppa NE (2015) Metabotropic glutamate receptors promote disinhibition of olfactory bulb glomeruli that scales with input strength. J Neurophysiol 113:1907–1920.

